# Root twisting drives halotropism via stress-induced microtubule reorientation

**DOI:** 10.1101/2022.06.05.494861

**Authors:** Bo Yu, Wenna Zheng, Lu Xing, Jian-Kang Zhu, Staffan Persson, Yang Zhao

## Abstract

Plants have evolved signaling mechanisms that guide growth away from adverse environments that can cause yield losses. Root halotropism is a sodium-specific negative tropism that is crucial for surviving and thriving under high salinity. Although root halotropism was discovered some years ago, the underlying molecular and cellular mechanisms remain unknown. Here, we show that abscisic acid (ABA)-mediated root twisting determines halotropism in *Arabidopsis*. An ABA-activated SnRK2 protein kinase (SnRK2.6) phosphorylates the microtubule-associated protein SP2L at Ser406, which induces a change in the anisotropic cell expansion at the root transition zone that located between the apical meristem and basal elongation zone, and is required for root twisting during halotropsim. Salt stress triggers SP2L-mediated cortical microtubule reorientation in cells at the transition zone, which guides cellulose microfibril patterns. Our findings outline the cellular and molecular mechanisms of root halotropism and indicate that anisotropic cell expansion through MT-reorientation and microfibril deposition have a central role in mediating tropic responses.

## INTRODUCTION

As sessile organisms, plants have evolved tropic and nastic movements that help them grow towards an optimal environment, which is crucial for plants to avoid deleterious microenvironments and analogous to the movement of animals. Plant organs thus integrate environmental signals to optimize growth, including light (phototropism), gravity (gravitropism), nutrients (nutritropism), chemicals (chemotropism), contact (thigmotropism), water (hydrotropism), and salinity (halotropism) (Gupta et al., 2020; Oldroyd and Leyser, 2020; Qu et al., 2015; Rosquete and Kleine-Vehn, 2013). Although root halotropism has been defined recently and is known as a response to avoid excessive Na^+^ but not Cl^-^ (Galvan-Ampudia et al., 2013; Sun et al., 2008), the underlying cellular and molecular mechanisms remain elusive.

Halotropism, i.e., a sodium-specific negative tropism, repels root grow from saline conditions, and is related to Na^+^ expulsion and Na^+^/K^+^ homeostasis (Deolu-Ajayi et al., 2019; Galvan-Ampudia et al., 2013). Salinity may be perceived by Glycosyl Inositol Phosphoryl Ceramides (GIPCs) at the cell-surface, or by sensing salt-induced cell wall changes (Feng et al., 2018; Jiang et al., 2019; Zhao et al., 2018a). These cues may trigger cytosolic Ca^2+^ release, sensed by the EF-hand Ca^2+^-binding protein Salt Overly Sensitive3 (SOS3, also named CBL4). SOS3 interacts with, and activates, the SOS2 protein kinase, which in turn phosphorylates the Na^+^/H^+^ antiporter SOS1, activating it to export excess Na^+^ ions out of cells (Guo et al., 2001). However, it remains unclear if these salt signaling components regulate halotropism.

High salinity may also cause hyperosmotic stress, which triggers hydrotropism (Eapen et al., 2005; Jaffe et al., 1985). Hydrotropism depends on MIZU-KUSSEI1 (MIZ1) and the plant hormone signaling mediated by abscisic acid (ABA) and cytokinins (Antoni et al., 2013; Chang et al., 2019; Kobayashi et al., 2007). MIZ1 is expressed in cortical cells of the root elongation zone and is localized in the cytoplasm and endoplasmic reticulum (ER) (Yamazaki et al., 2012). MIZ1 inhibits the activity of ER Ca^2+^-ATPase isoform ECA1 and mediates a slow, long-distance, phloem-transmitted, asymmetric Ca^2+^ signal in the elongation zone (Shkolnik et al., 2018). ABA activates the plasma membrane H^+^-ATPase 2 (AHA2) by promoting its phosphorylation, which mediates an asymmetric H^+^ efflux that is crucial for differential growth of the elongation zone (Dietrich et al., 2017; Miao et al., 2021). MIZ1 is also expressed in the root meristem, where it mediates asymmetric cytokinin responses (Chang et al., 2019; Yamazaki et al., 2012). Thus, MIZ1 may regulate hydrotropism by controlling Ca^2+^ signals and hormone homeostasis.

Auxin distribution is key to asymmetric plant growth (Robert and Friml, 2009). Notably, asymmetric auxin distribution occurs two hours after halo-stimulation in roots, and is thought to drive the halotropic root bending, fitting the widely accepted Cholodny-Went theory (Galvan-Ampudia et al., 2013; Went and Thimann, 1937). Nevertheless, endocytosis of key auxin efflux carrier PIN-FORMED (PIN) 2 did not occur until six hours after halo-stimulation (Galvan-Ampudia et al., 2013). While not required during the gravitropic response, PIN2 endocytosis somewhat dampens asymmetric auxin distribution (Retzer et al., 2019). Because gravitropism antagonizes halotropic root bending, phospholipase D (PLD) ζ1 and ζ2 may be involved in halotropism by countering gravitropism (Galvan-Ampudia et al., 2013; Korver et al., 2020; McLoughlin et al., 2013; Sun et al., 2008).

Plant cell walls encase plant cells. The mechanical inhomogeneity of the cell wall, the cell wall and plasma membrane connection, and vacuolar expansion regulate anisotropic growth and thus plant morphology (Chebli et al., 2021; Du et al., 2022; Dunser et al., 2019; Gorelova et al., 2021; Lin et al., 2022; Tang et al., 2022; Zhao et al., 2020). Cellulose, produced at the cell surface by cellulose synthases (CESAs) is the load-bearing structure of cell walls (Chebli and Geitmann, 2017). Notably, cortical microtubules (MTs) direct how cellulose is organized and thus growth anisotropy (Endler et al., 2015; Gu et al., 2010). Many MT-associated proteins (MAPs) influence MT organization, including the MT-severing enzyme KATANIN that critically redirect MT organization during seedling de-etiolation (Lindeboom et al., 2013). Another important MAP is SPIRAL2 (SPR2, also known as TORTIFOLIA1/TOR1), characterized by multiple N-terminal HEAT repeats, which together with its C-terminus, aid in MT interactions (Buschmann et al., 2004; Shoji et al., 2004; Yao et al., 2008). Whereas *SPR2* is strongly expressed in various tissues, its close homolog *SP2L* is weakly expressed but relatively abundant in the hydathodes and roots (Yao et al., 2008). SPR2 binds MT plus- and minus-ends and MT intersections (Fan et al., 2018; Leong et al., 2018; Nakamura et al., 2018; Wightman et al., 2013) and is the only known protein to track with and stabilize the treadmilling MT minus ends in land plants (Fan et al., 2018; Leong et al., 2018; Nakamura et al., 2018). SP2L and SPR2 have overlapping functions and regulate MT-end dynamics and anisotropic organ growth (Buschmann et al., 2004; Shoji et al., 2004; Yao et al., 2008). Moreover, SPR2 functions together with KATANIN during blue light-induced MT array reorientation in elongating hypocotyl cells to promote MT severing at crossovers (Fan et al., 2018; Lindeboom et al., 2013; Nakamura et al., 2018). Ectopic expression of *SP2L* can complement the phenotype of *spr2* mutant (Yao et al., 2008), indicating that SP2L may have similar functions as SPR2, e.g., in mediating MT array reorientation.

We show that root halotropism does not require MIZ1 protein function and does not depend on the functional cytokinin signaling. Although auxin redistribution may not be necessary for the early stage of halotropic root bending, it plays an essential role during the later stage, probably by resuming gravitropism. Instead, we demonstrate that the ABA-activated protein kinase SnRK2.6 phosphorylates a single residue on SP2L, which drives cortical MT reorientation at the root transition zone that located between the apical meristem and basal elongation zone (Baluška et al., 2010; Kong et al., 2018), to change growth anisotropy and thus halotropism. Thus, we present a Cholodny-Went theory-independent mechanism in which stimuli-induced MT reorientation determines tropic growth during root halotropism in *Arabidopsis*.

## RESULTS

### Root halotropism does not require classical salt stress signaling

To investigate molecular mechanisms that underpin root halotropism in *Arabidopsis*, we optimized the vertical split-agar assay in which root tips were subjected to diagonal NaCl gradients (Galvan-Ampudia et al., 2013) (Figure 1A). Roots changed their growth direction away from the high salinity region when seedlings were transferred to the split-agar system containing 200 mM NaCl, but grew downward following the gravity axis on split-agar control medium (Figures 1B and 1C). To first investigate whether classical salt signaling mediates halotropism, we analyzed the halotropic root bending in mutants defected in sensing or signaling of salt stress. These included *moca1*, defective in GIPC contents that impact Ca^2+^ signals (Jiang et al., 2019), and *sos* mutants, which are deficient in Ca^2+^-mediated Na^+^-transport (Yang and Guo, 2018). Both the *sos* and *moca1* mutants are hypersensitive to salt stress (Yang and Guo, 2018). Interestingly, halotropic root bending was enhanced in both of these mutants (Figure 1D & E). Since *sos* and *moca1* mutants accumulate excessive Na^+^, we speculate that halotropism is triggered by ionic stress of the root tips. We next analyzed Na^+^ accumulation in root tips using the Na^+^-specific fluorescent indicator CoroNa Green (Figure 1F) (Meier et al., 2006; Park et al., 2009). As expected, Na^+^ accumulation was significantly higher in the *moca1, sos2* and *3* mutants as compared to wild-type root tips (Figures 1F and 1G). We propose that the enhanced halotropism in *sos* and *moca1* mutants is linked to Na^+^ accumulation in the root tips and that the halotropic response is controlled by a hitherto unknown pathway, rather than via classical salt stress signaling.

**Figure 1.**
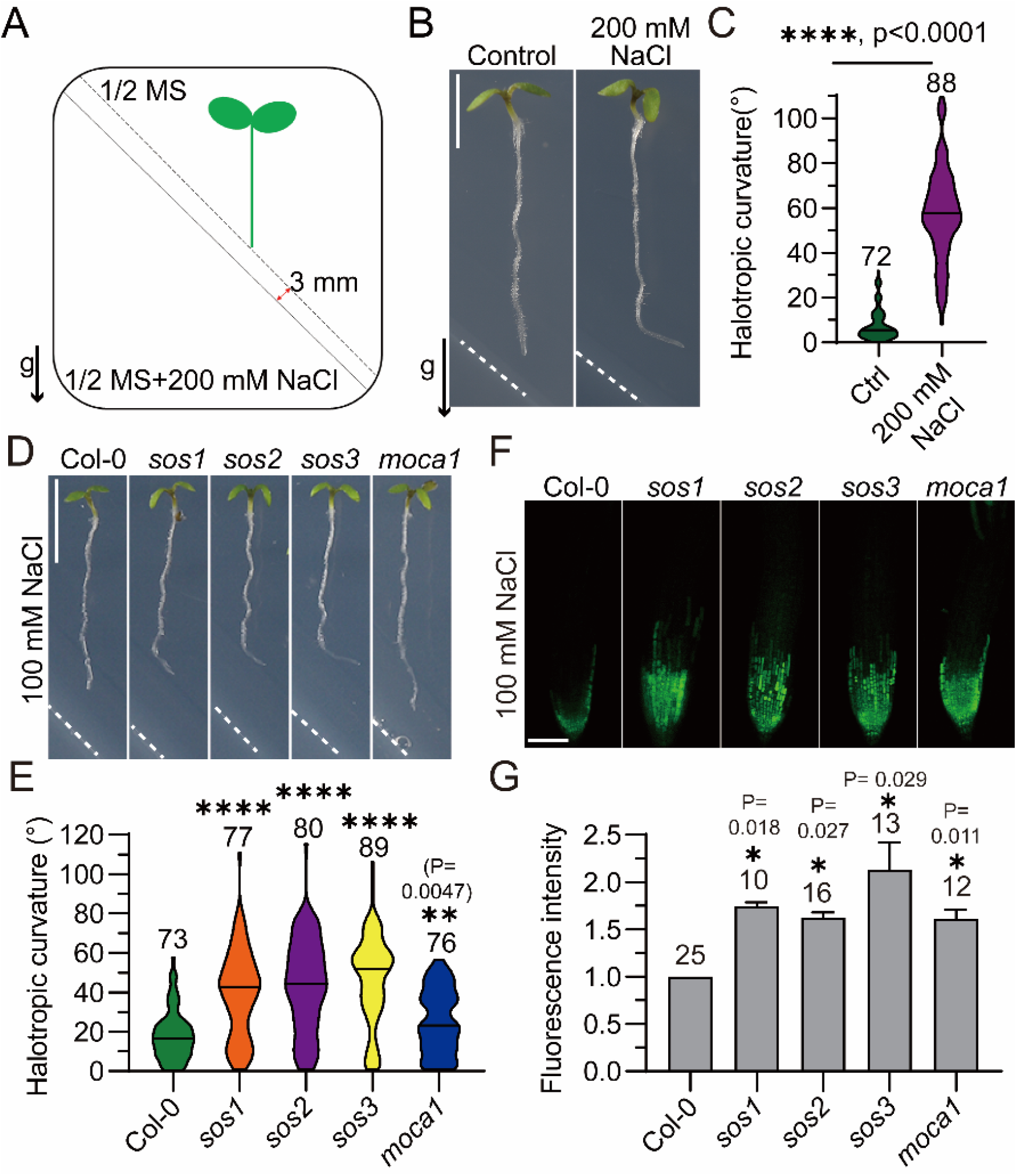
Root halotropism does not require classical salt-stress signaling. (A) Schematic diagram of halotropism assay. Four-day-old seedlings with straight roots were transferred from 1/2 MS vertical plates to 1/2 MS split-agar medium containing 200 mM NaCl at the bottom left side. Root tips were 3 mm above the boundary of the two agar mediums. (B-C) Wild-type (Col-0) seedlings were transferred to a split-agar medium without or with 200 mM NaCl. Scale bar, 0.5 cm. (B). The halotropic root curvature was quantified (C). 0° equals vertical. Values are mean ± s.e.m (n = 72-88 seedlings). (D-E) Halotropic root bending of *sos* and *moca1* mutants on split-agar medium with 100 mM NaCl (D). Scale bar, 0.5 cm. The halotropic root curvature was quantified (E). Values are mean ± s.e.m (n = 15-36 seedlings). (F-G) Seedlings were stained with CoroNa Green fluorescent dye. Seedlings were imaged after 3 hours of halo-stimulation (F). Scale bar, 100 μm. Fluorescent intensities in root tips were quantified (G). Values are mean ± s.e.m (n = 10-25 seedlings).

### MIZ1, auxin distribution, and cytokinin signaling do not control root halotropism

Asymmetric growth of organs is mainly controlled by asymmetric distribution of auxin or cytokinins (Chang et al., 2019; Su et al., 2017). While asymmetric auxin distribution occurs during halotropic root tip responses (Galvan-Ampudia et al., 2013), its requirement for *Arabidopsis* root halotropism remains unclear. In roots, auxin is distributed by PIN auxin efflux carriers and AUXIN1/LIKE-AUX1 (AUX1/LAX) auxin influx carriers (Su et al., 2017). To our surprise, halotropic root bending was not blocked, but instead enhanced, in *aux* or *pin* mutants, especially in *aux1* and *pin2*, which are both impaired in gravitropic responses (Figures 2A, S1A and S1B). In contrast, the *pin2* mutant roots were randomly oriented after transferring to a control medium in split agar assays, with an average value similar to WT (Figure S1C). Since the halotropic root bending has to challenge gravity, the enhanced root bending in *aux1* and *pin2* could be caused by the impaired gravitropism but not enhanced halotropism. YUCCA (YUC) enzymes are rate-limiting in the main auxin biosynthesis pathway in *Arabidopsis* (Zhao, 2018). Notably, halotropic root bending was similar to wild type in the *yuc* mutants (Figures 2A, 2B, S1A and S1B). While we cannot rule out that certain aspects of halotropic responses are related to auxin biology, we conclude that halotropic root bending may not depend on the canonical auxin synthesis and redistribution framework.

**Figure 2.**
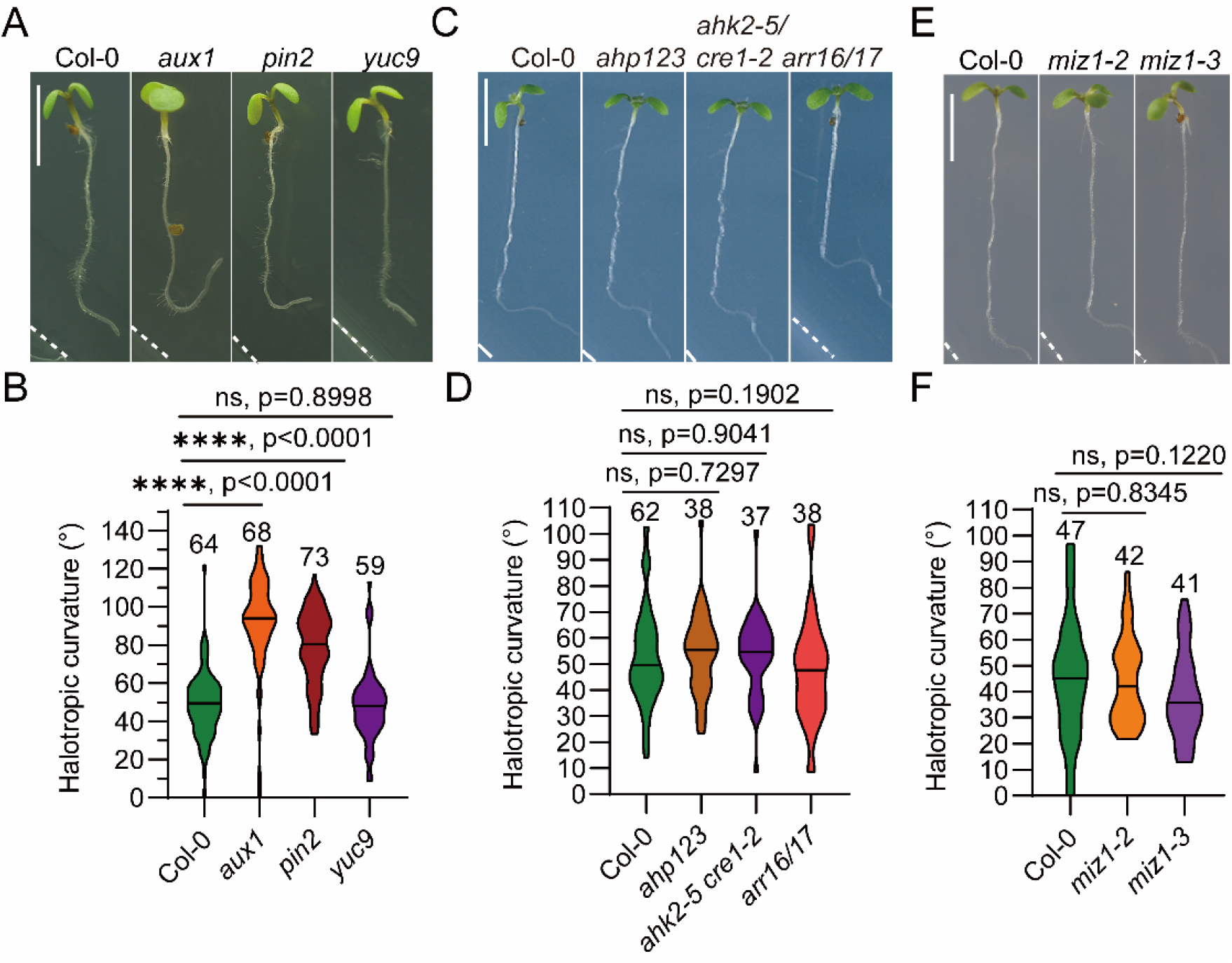
MIZ1, auxin redistribution, and cytokinin signaling do not explain root halotropism. Wild-type (Col-0), auxin-related mutants (A-B), cytokinin signaling mutants (C-D), and hydrotropism defective *miz1* mutants (E-F) were transferred to split-agar medium with 200 mM NaCl. The halotropic root curvature was quantified (B, D, and F). Values are mean ± s.e.m (n = 37-73 seedlings).

Asymmetric cytokinin response is required for root hydrotropism (Chang et al., 2019). High salinity causes hyperosmotic stress that triggers hydrotropism. Therefore, we analyzed root halotropism in hydrotropism-defective, cytokinin-related mutants, including *ahk2-5/cre1-2, ahp1/2/3*, and *arr16/arr17* (Chang et al., 2019). These mutants exhibited similar halotropic responses as wild-type seedlings (Figures 2C and 2D). Moreover, the central regulator of root hydrotropism, *MIZ1*, did not contribute to root halotropism, as *miz1* mutants responded similar to wild type (Figures 2E and 2F, S1D and S1E). These results indicate that root halotropism is controlled by different mechanisms than hydrotropism and may be considered a novel tropism.

### Root halotropism requires ABA production and signaling

ABA is central to plant abiotic stress responses, including salt, and might thus regulate halotropic root bending. To test this, we used ABACUS, a FRET sensor for ABA, to measure ABA levels (Jones et al., 2014). As expected, halo-stimulation significantly increased ABA accumulation in root tips within two hours after treatment (Figure 3A and 3B). In the *nced3/5* mutant, stress-induced ABA synthesis is reduced (Frey et al., 2012), while the ABA signal is almost abolished in the *snrk2*.*2/3/6* and the *pyl* duodecuple mutants (Fujii and Zhu, 2009; Zhao et al., 2018b). We used these three mutants to test whether ABA is necessary for root halotropism. We found that root halotropic bending was significantly reduced in the *nced3/5, snrk2*.*2/3/6*, and *pyl* duodecuple mutants (Figures 3C and 3D), indicating that ABA signaling is important for root halotropism.

**Figure 3.**
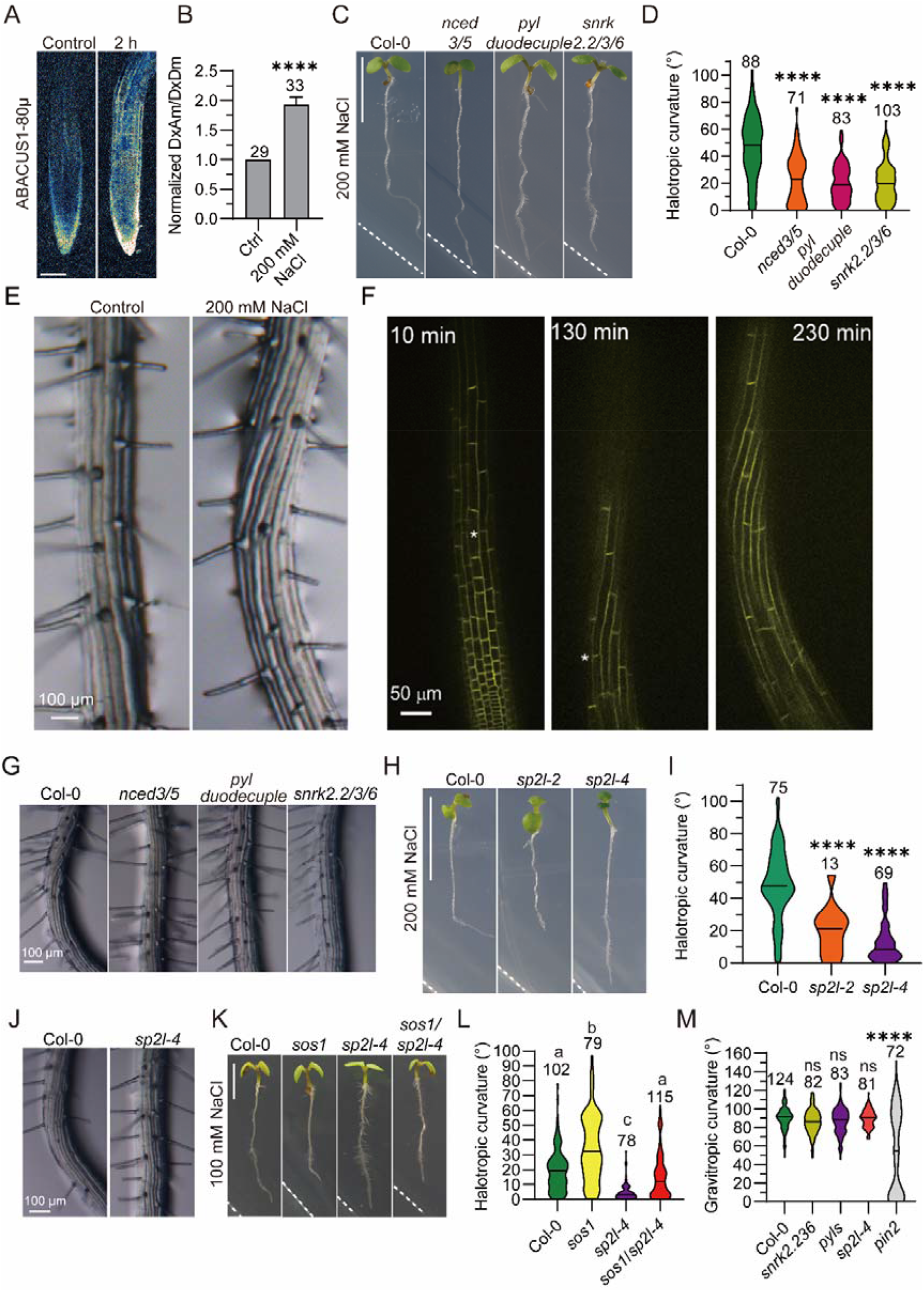
ABA and anisotropic cell expansion are crucial for root halotropism. (A-B) ABA accumulation in root tips using ABACUS1-80μ without or with halo-stimulation on split-agar medium. Scale bar, 100 μm. ABA accumulation was quantified (B). Values are mean ± s.e.m (n = 29-33 seedlings). (C-D) Wild-type (Col-0), ABA biosynthesis (*nced3/5*), and signaling mutants (*pyl* duodecuple and *snrk2*.*2/3/6*) were transferred to split-agar medium with 200 mM NaCl (C). The halotropic root curvature was quantified (D). Scale bar, 0.5 cm. Values are mean ± s.e.m (n = 71-103 seedlings). (E-F) Anisotropic cell expansion in root epidermal of the bending region. Root growth was monitored using Col-0 (wild-type) under a stereoscopic microscope 18 h after halo-stimulation (E), or a YFP plasma membrane reporter *NPSN12* line for observation of cell boundaries under a Nikon-Andor WD spinning disc confocal with vertical stage 10-230 min after halo-stimulation (F). Scale bars, 100 μm in (E), and 500 μm in (F). (G) Root twisting in the mature zone is monitored under a stereoscopic microscope 18 h after halo-stimulation. Scale bar, 100 μm. (H-J) Wild-type (Col-0), and *sp2l* mutants were transferred to split-agar medium with 200 mM NaCl (H). The halotropic root curvature was quantified (I). Root growth is monitored under a stereoscopic microscope (J). Scale bars, 0.5 cm in (H), and 100 μm in (I).Values are mean ± s.e.m (n = 13-75 seedlings). (K-L) Wild-type (Col-0), *sos1, sp2l-4* mutants, and *sos1/sp2l-4* mutants were transferred to split-agar medium with 100 mM NaCl (K). The halotropic root curvature was quantified (L). Scale bar, 0.5 cm. Values are mean ± s.e.m (n = 6 seedlings), with different letters (a and b) indicating significant difference as evaluated by post-hoc Tukey test after ANOVA (P < 0.05). (M) Root gravitropic growth of ABA-insensitive mutants (*pyl* duodecuple and *snrk2*.*2/3/*6) and the *sp2l-4* mutant, with pin2 and Col-0 as controls. The gravitropic root curvature was quantified. Values are mean ± s.e.m (n = 72-124 seedlings). ** *P* <0.01, **** *P* <0.0001, Student’s t-test.

### Root halotropism requires anisotropic cell expansion

Changes in growth patterns often involve microtubule re-organization (Chebli et al., 2021; Furutani et al., 2000). To first assess if halotropic root bending is due to changes in root cell expansion, we undertook a detailed analysis of root growth and cell morphology during this process. Changes in root growth directions were observed as early as two hours after halo-stimulation (Figures 3E, S2A and S2B). To better discern changes in cell morphology, we used the EYFP-NPSN12 marker that outlines the plasma membrane in elongating epidermal cells of seedling roots using a Nikon-Andor WD spinning disc confocal microscope with a vertical stage (Figures 3F and S2C). Interestingly, the root epidermal cells exhibited a right-handed twist within the bending region of roots (Figure 3E and 3F). Moreover, anisotropic cell expansion changed in the epidermis of root transition zone that located between the apical meristem and basal elongation zone, and was restricted to the bending region of the root (Figures 3F, S2B and S2C). The root epidermal cell twist did not occur in the corresponding region of roots of ABA biosynthesis and signaling mutants (Figure 3G), indicating that ABA might regulate root halotropism by inducing changes in root cell growth patterns.

Anisotropic cell expansion is tightly controlled by cytoskeletal arrangements, which in turn are regulated by multiple MT-associated and actin-binding proteins (Buschmann et al., 2004; Chen et al., 2016; Shoji et al., 2004; Wang and Mao, 2019). Given that ABA signaling is necessary for halotropic root bending, we hypothesized that key ABA signaling components may drive cytoskeletal re-arrangements during this process. We paid particular attention to SnRK2 protein kinases due to the block in halotropic root bending in the *snrk2*.*2/3/6* mutant. We screened the literature for putative SnRK2-mediated cytoskeleton-related processes. Interestingly, several cytoskeleton-related proteins are putative SnRK2 substrates according to phosphoproteomic analysis in the *snrk2*.*2/3/6* triple mutant (Figure S2D) (Umezawa et al., 2013; Wang et al., 2013). We next conducted a reverse genetic screen based on root halotropic response phenotypes using mutants of these potential SnRK2 substrates (Figure S2E). Here, we identified the MAP SP2L as a component required for root halotropism. Indeed, two T-DNA insertion knockout mutants, *sp2l-2* and SAIL_117_F12 (named *sp2l-4*) (Figure S2F) (Yao et al., 2008), were defective in root halotropism and, consequently, did not display the right-handed twist of elongating root cells (Figures 3H to 3J). To explore interactions between *sp2l* and hypersensitive halotropic growth mutants, we crossed *sp2l-4* with *sos1*. The resulting double mutant did not respond to halo-stimulation, thus mimicking the *sp2l* mutant phenotype and indicating that SP2L acts downstream of SOS1 (Figures 3K and 3L). Since gravitropism antagonizes halotropic root bending (Figures 2A, 2B, S1A and S1B), we next analyzed root gravitropic growth of the ABA-insensitive and *sp2l* mutants. The ABA-insensitive and *sp2l-4* mutants exhibited similar root curvature as wild-type seedlings during gravitropic root growth (Figure 3M), indicating that ABA signaling and SP2L are not required for gravitropism. These results demonstrated that ABA and SP2L act downstream of salt-stress sensing.

### SnRK2.6 interacts with and phosphorylates SP2L

The SP2L protein contains multiple HEAT repeats, which serve as MT-binding domains at its N-terminus, and a MT-binding region in its C-terminus (Figure 4A). To test potential interactions between SP2L and SnRK2.6, we fused SP2L to the GAL4 DNA-binding domain (BD) and SnRK2.6 to the GAL4-activating domain (AD), performed yeast two-hybrid assays, and detected interactions between SP2L and SnRK2.6 (Figure 4B). We confirmed this interaction using bimolecular fluorescent complementation (BiFC) in *Nicotiana benthamiana* leaves (Figure 4C). Because the full-length SP2L proteins are difficult to purify and are prone to degradation, we purified a truncated SP2L protein, SP2L-MC (middle region and C-terminal, Figure 4A). The ST- and HIS-tagged SnRK2.6 protein interacted with GST-SP2L-MC-GFP but not with the negative control GST-GFP (Figure 4D) in pull-down assays. Therefore, we concluded that SP2L interacts with SnRK2.6.

**Figure 4.**
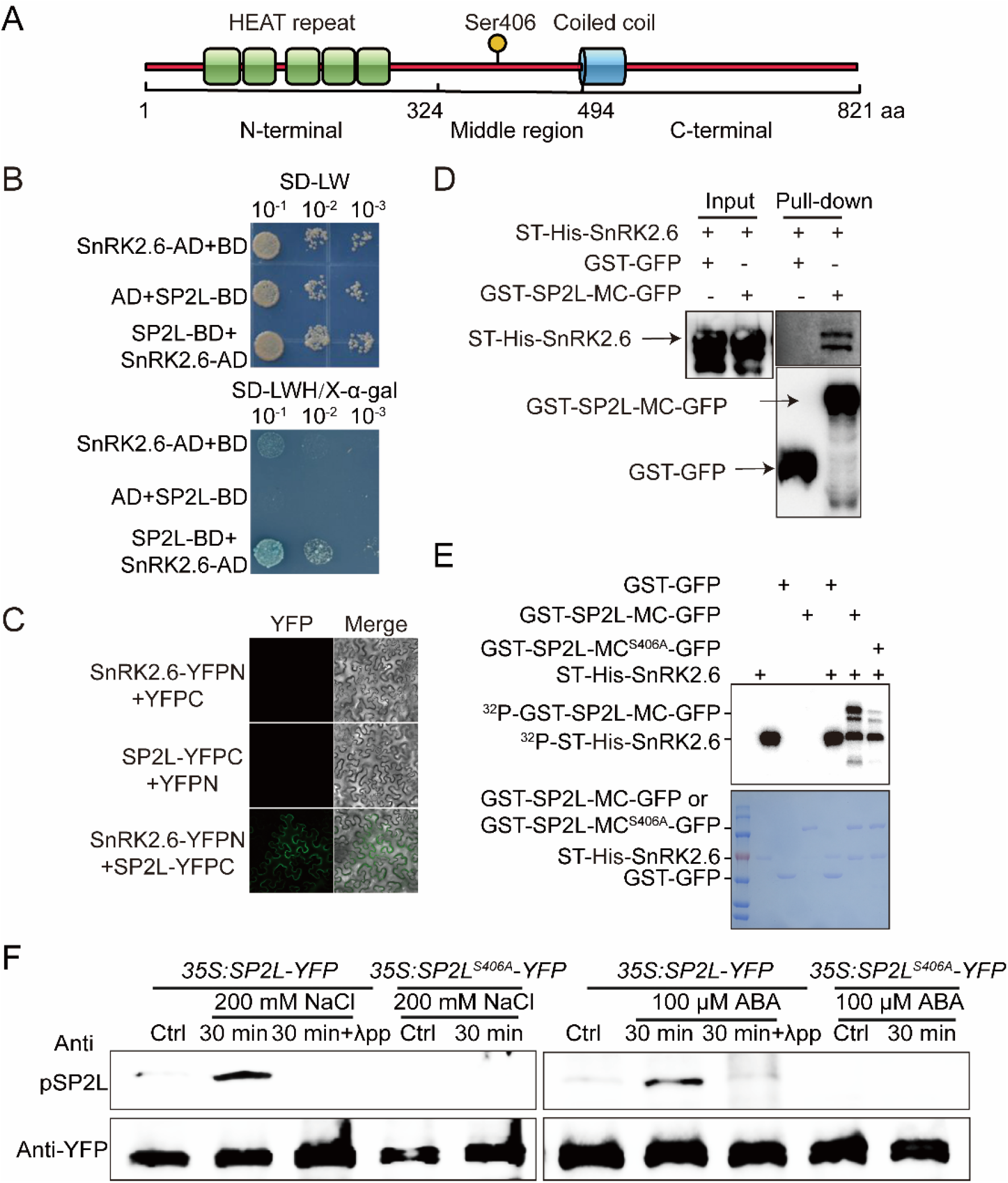
SnRK2.6 interacts with and phosphorylates SP2L. (A) Protein domains of SP2L. The SP2L protein has five HEAT repeats that are predicted to bind MT at the N-terminus, and a coiled-coil domain in the MT-binding C-terminal region. The phosphorylation site Ser406 is located in the middle region. (B) SnRK2.6-SP2L interaction in yeast two-hybrid assay. Interactions were evaluated by yeast growth on media lacking Leu, Trp, and His (SD-LWH) supplemented with X-α-Gal (20 mg/ml). Yeast was diluted 10 times (10^−1^), 100 times (10^−2^), and 1000 times (10^−3^). Combinations of SnRK2.6-AD with BD, and SP2L-BD with AD were used as negative controls. (C) Interactions between SnRK2.6 and SP2L in *N. benthamiana* leaves using bimolecular fluorescence complementation assays (BiFC). Full-length SnRK2.6 and SP2L were fused to the split N- or C-terminal fragments of YFP (SnRK2.6-YFPN and SP2L-YFPC). Unfused YFPN and YFPC were used as negative controls. Merge: merged images of YFP channel and brightfield. (D) Interactions between SnRK2.6 and SP2L-MC in a pull-down assay. Interactions were determined by the co-immunoprecipitation of ST-His-SnRK2.6 with GST-SP2L-MC-GFP protein using Glutathione Sepharose beads. Immunoblot analyses were detected with anti-His antibody and anti-GST antibody. GST-GFP protein was used as a negative control. (e) Phosphorylation of SP2L-MC and SP2L-MC-S406A by recombinant SnRK2.6. ST-His-SnRK2.6 was incubated with wild-type or mutated SP2L-MC in protein kinase buffer containing [γ-^32^P] ATP. Protein phosphorylation and loading were detected by autoradiography (top panel) and Coomassie staining (bottom panel) after gel electrophoresis. (F) Phosphorylation of Ser406 of SP2L under salt stress or ABA treatment in *35S:SP2L-YFP* and *35S:SP2L-S406A-YFP* transgenic plants. An anti-phospho-Ser406-SP2L antibody was used to detect phosphorylation of SP2L. Protein loading was detected by anti-YFP. λPP: Lambda protein phosphatase.

We next asked if SP2L is a substrate of SnRK2 phosphorylation. *In vitro* kinase assays showed that SnRK2.6 could phosphorylate truncated SP2L-MC proteins (Figure 4E). We performed mass spectrometry analyses to identify the potential phosphorylation sites and found that SnRK2.6 phosphorylates SP2L mainly at Ser406 (Figure S3A). We mutated the serine (S) to a similar-sized, non-phosphorylatable alanine (A). We found that the S406A mutation reduced the phosphorylation of the truncated SP2L-MC (Figures 4E).

To examine whether salt stress induces phosphorylation of SP2L *in vivo, SP2L-YFP*-expressing seedlings were treated with 200 mM NaCl for 30 min, which induced phosphorylation on Ser406 through IP-MS assays (Figure S3B). To confirm this result, we generated antibodies that recognize Ser406-phosphorylated SP2L using a modified peptide KLEKRG-pS-GD as antigen. The anti Ser406 phosphorylation antibody specifically recognized phosphorylated Ser406 on SP2L-MC after *in vitro* kinase assays (Figure S3C). To verify the salt stress-induced phosphorylation of Ser406, we generated *35S:SP2L-YFP* and *35S:SP2L-S406A-YFP* transgenic lines and performed immunoblot analysis using the anti-Ser406 phosphorylation antibody. Salt stress and ABA treatment both induced phosphorylation of Ser406 in SP2L, while this phosphorylation was not detected in the mutated protein (Figure 4F). The phosphorylation was effectively removed by λ-protein phosphatase (λ-PPase) treatment (Figures 4F). These results strongly support that salt stress and ABA induce phosphorylation of Ser406 in SP2L via SnRK2.

### Serine 406 phosphorylation of SP2L is essential for root halotropism

To test the biological significance of S406 phosphorylation on SP2L in root halotropism, we generated transgenic lines with native promoter-driven constructs of either a wild-type *SP2L* or a non-phosphorylatable *SP2L-S406A* in the *sp2l-4* background (Figures 5A and 5B). These transgenic lines have a similar expression level of *SP2L* as Col-0 wild-type (Figure 5C). The wild-type *SP2L* completely recovered the root halotropism defects of the *sp2l-4* mutant (Figures 5A and 5B). In contrast, expression of the *SP2L-S406A* did not rescue the root halotropism defects of the *sp2l-4* mutant (Figures 5A and 5B). Likewise, the lack of the right-handed twist in the root epidermal cells exposed to salt-containing medium was rescued by the wild-type *SP2L*, but not by the *SP2L-S406A* (Figure 5D). These results indicate that phosphorylation at serine 406 mediates root halotropism by inducing a change in anisotropic cell expansion.

**Figure 5.**
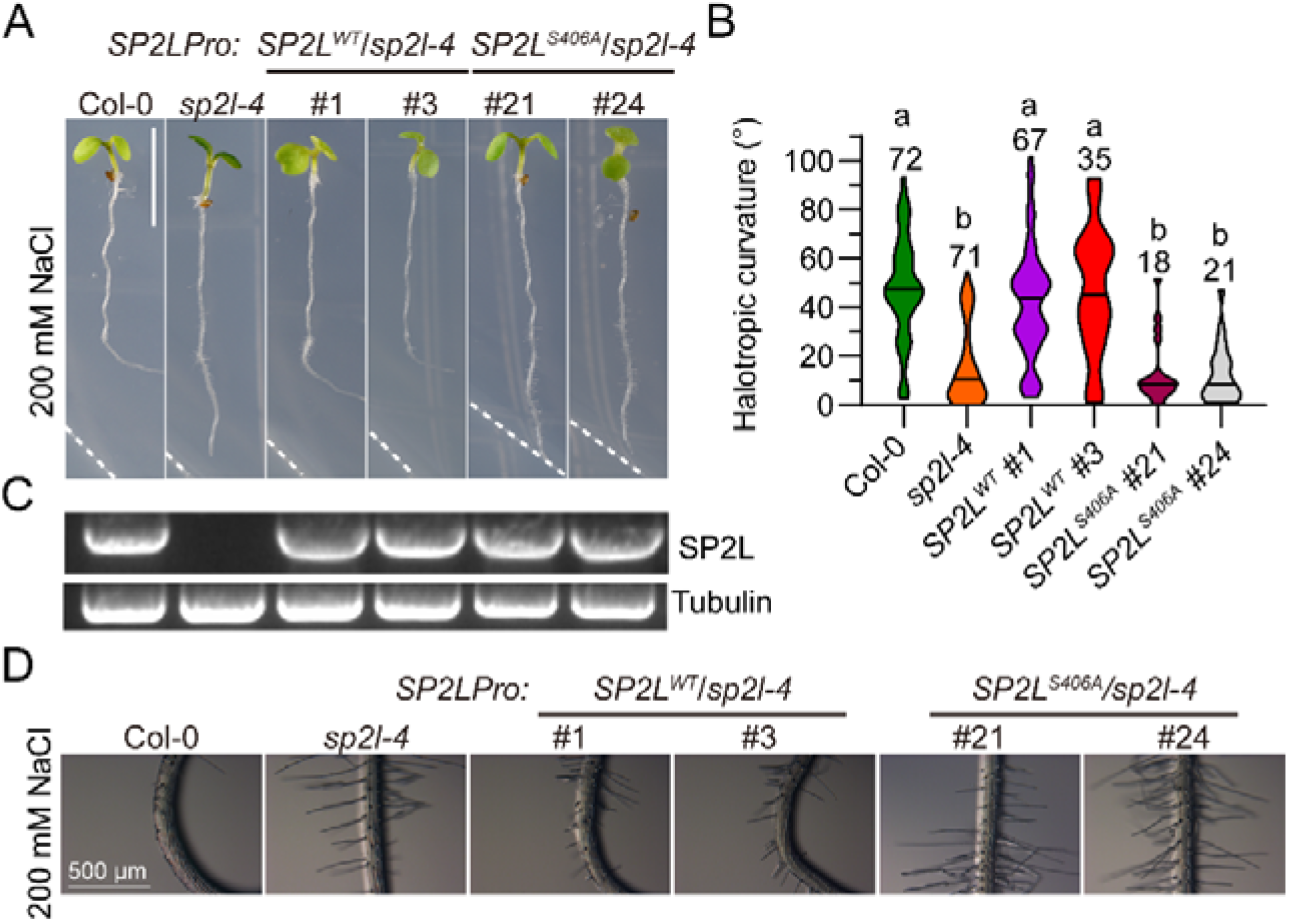
Phosphorylation of SP2L is essential for root halotropism. Root halotropic growth of Col-0 wild-type, *sp2l-4* mutant, and *SP2Lpro:SP2L* transgenic plants expressing wild-type or *S406A*-mutated *SP2L* in *sp2l-4* mutant background. The photo was taken 24 hours after transferring seedlings to a split-agar medium with 200 mM NaCl (A). The halotropic root curvature was quantified (B). Expression of *SP2L* in corresponding lines was analyzed by RT-PCR, and tubulin was used as an internal reference (C). Root growth was monitored under a stereoscopic microscope (D). Values are mean ± s.e.m (n = 18-72 seedlings). Scale bar, 0.5 cm in (A), and 500 μm in (D).

### SP2L promotes MT reorientation upon halo-stimulation in elongating root cells

SP2L-GFP associates with cortical MT arrays in root and hypocotyl epidermal cells when stably expressed under the control of the native promoter (Yao et al., 2008). In our hands, we only observed very weak fluorescence with native promoter driven fluorescent constructs. To confirm the localization, we instead used *35S:SP2L-mCherry* and transformed this construct into transgenic lines harboring the MT-marker *35S:GFP-TUB6*. The SP2L-mCherry signal extensively overlapped with cortical MTs labeled by GFP-TUB6 in hypocotyl epidermal cells (Figures 6A and 6B). Thus, we studied the role of SP2L in cortical MT array reorientation during root halotropism.

**Figure 6.**
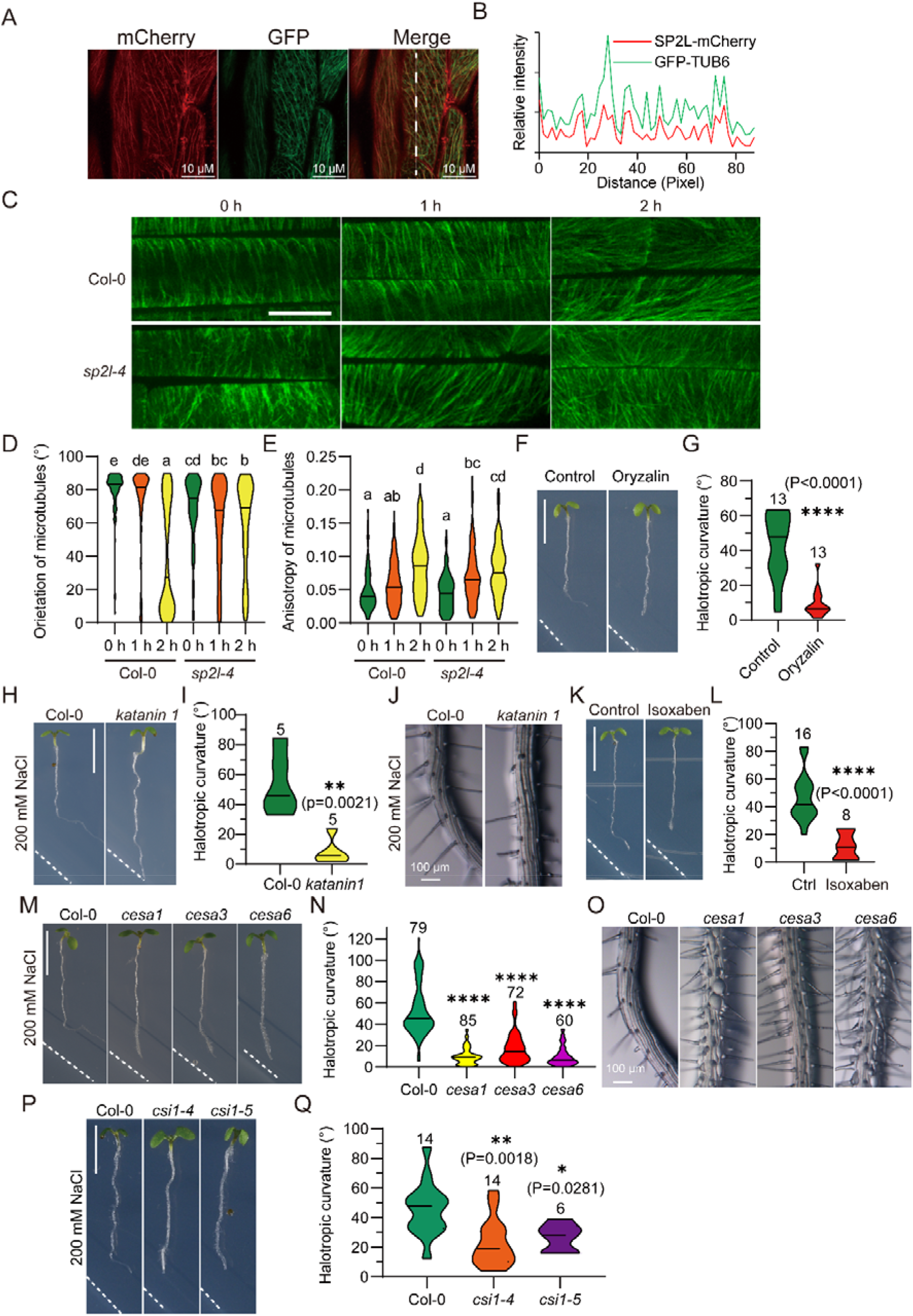
SP2L promotes microtubule reorganization upon halo-stimulation. (A-B) SP2L-mCherry (left) and GFP-TUB6 labeled MTs (middle) in hypocotyl epidermal cells. The merged image is shown on the right (A). Scale bars, 10 μm. The fluorescence intensities (mCherry and GFP signals) were scanned (B). (C) Images of cortical microtubules in WT and *sp2l-4* root epidermal cells in the transition zone after halostimulation, as visualized by expression of GFP-TUB. Scale bar, 10 μm. (D-E) Quantification of cortical MT orientation and anisotropy in (C). Data are mean values from 18-20 independent cells. (F-G) Root halotropic growth of Col-0 wild-type without (left panel, control) or with (right panel) 1 μM oryzalin treatment. The photo was taken 24 hours after transferring seedlings to a split-agar medium with 200 mM NaCl (F). Scale bar, 0.5 cm. The halotropic root curvature was quantified (G). Values are mean ± s.e.m (n = 13). (H-J) Root halotropic growth of Col-0 wild-type and *katanin1* mutant. The photo was taken 24 hours after transferring seedlings to a split-agar medium with 200 mM NaCl (H). Scale bars, 0.5 cm in (H), and 100 μm in (J). The halotropic root curvature was quantified (I). Root growth was monitored under a stereoscopic microscope (J). Values are mean ± s.e.m (n = 5). (K-L) Root halotropic growth of Col-0 wild-type without (left panel, control) or with (right panel) 1 nM isoxaben treatment. The photo was taken 24 hours after transferring seedlings to a split-agar medium with 200 mM NaCl (K). Scale bar, 0.5 cm. The halotropic root curvature was quantified (L). Values are mean ± s.e.m (n = 8-16). (M-Q) Root halotropic growth of Col-0 wild-type, *cesa1* (*rsw1-1*), *cesa3* (*ixr1-2*), and *cesa6* (*prc1-1*) mutants, and *csi1* mutants. The photo was taken 16 hours after transferring seedlings to a split-agar medium with 200 mM NaCl (M and P). Scale bars, 0.5 cm in (M and P), and 100 μm in (O). The halotropic root curvature was quantified (N and Q). Root growth was monitored under a stereoscopic microscope (O). Values are mean ± s.e.m (n = 32-34).

Since the change in anisotropic cell expansion happens in the root transition zone as early as two hours after halo-stimulation (Figure S2A and S2C), we imaged GFP-TUB-labeled cortical MTs in the elongating epidermal cells at the root transition zone within this period. Initially, the cortical MTs were transversely oriented and, approximately two hours after halo-stimulation, were reoriented to oblique, or even longitudinal, direction (Figure 6C and 6D). In contrast, cortical MT reorientation in the elongating epidermal cells did not respond in the same way in the *sp2l-4* mutant after halo-stimulation (Figure 6C and 6D). The anisotropy of cortical MTs was increased slightly both in Col-0 wild-type and *sp2l-4* mutant (Figure 6E). In addition, treatment with the MT depolymerizing agent oryzalin blocked root halotropic response (Figures 6F and 6G), supporting our hypothesis that MT organization is critical for halotropism, although we could not avoid side effects caused by the chemical. These results indicated that SP2L functions in mediating cortical MT reorientation in the elongating epidermal cells during root halotropism.

Blue light-induced cortical MT reorientation is controlled by KATANIN-mediated severing at MT crossovers to generate a new MT organization (Lindeboom et al., 2013). This process is supported by SPR2-mediated MT minus-end stabilization that increases MT crossover lifetime and severing (Fan et al., 2018; Lindeboom et al., 2013; Nakamura et al., 2018). Therefore, we speculated that SP2L might function together with KATANIN in controlling root halotropism. We therefore assayed the root halotropic responses in *ktn1* mutants, corresponding to the catalytic KATANIN P60 subunit. Indeed, the *ktn1* mutant was defective in root halotropism (Figures 6H and 6I). Moreover, the salt-induced right-handed twist in the root epidermal cells was abolished in the *ktn1* mutant (Figure 6J). These results confirmed the regulation of root halotropism by MT reorientation.

### MT reorientation controls cellulose synthase patterns upon halo-stimulation

The rearrangement of cortical MTs redirects trajectories of CESA complexes to alter cellulose microfibril patterns through the MT-associated COMPANION OF CESA (CC) proteins and CESA INTERACTING1 (CSI1) (Endler et al., 2015; Gu et al., 2010). To study the CESA behaviour, we imaged GFP-labeled CESA3 in the elongating epidermal cells at the root transition zone in dual-labeled GFP-CESA3 and mCherry-TUA5 lines (Figure S4). The orientation of GFP-CESA3 changed from transverse to longitudinal orientation after two hours of halo-stimulation, similar to the reorientation of cortical MTs (Figure S4). Several CESAs, including CESA1, CESA3, and CESA6, are required for primary wall synthesis and anisotropic cell expansion (Persson et al., 2007). Indeed, pharmacologically disrupting cellulose biosynthesis, using the cellulose synthesis inhibitor isoxaben, impaired the root halotropic response (Figures 6K and 6L). To corroborate these findings, we assayed the root halotropic responses in *cesa1/rsw1-1, cesa3/ixr1-2, cesa6/prc1-1*, and *csi1/pom2* mutants. All three *cesa* mutants and the *csi1* mutants were defective in root halotropism (Figures 6M-6Q). In addition, the right-handed twist in the root epidermal cells was impaired in the *cesa* mutants (Figure 6O). These results suggested that SP2L-mediated MT-reorientation controls root halotropism by affecting cellulose microfibril deposition that drives right-hand twisting of elongating root cells.

### Root halotropism contributes to salt stress tolerance

Root halotropism helps roots grow away from a high salinity microenvironment. To investigate whether halotropism contributes to salt stress tolerance, we transferred wild-type, *pyl* duodecuple, *snrk2*.*2/3/6*, and *sp2l-4* mutant seedlings to split-agar medium with 200 mM NaCl. After plants were subjected to salt stress for 21 days, the ABA signaling mutants and *sp2l-4* mutants exhibited reduced shoot growth compared to wild-type (Figure S5A), possibly resulting from enhanced Na^+^ accumulation in the mutants, presumably caused by a relatively higher ratio of roots in the high salinity region. We further confirmed root halotropism using the vertical split-agar assay in which root tips were subjected to vertical NaCl gradients (Figure S5B). We found that both the primary and lateral roots undergo halotropic root bending (Figure S5C), suggesting that plants rearrange root architecture to survive and thrive under saline conditions.

## Discussion

High salinity is a major abiotic stress unevenly distributed in different soil layers dependent on precipitation, irrigation, and evaporation. Plants have evolved complex signaling systems to sense environmental factors and to guide them to grow away from adverse conditions (Lamers et al., 2020), which is essential for survival under changeable environments. In this work, we outlined the cellular and molecular mechanisms for root halotropism, in which ABA signaling, cortical MT reorientation, and microfibril deposition coordinate to mediate a right-handed change in anisotropic cell expansion in elongating root cells.

While salt stress tolerance and avoidance contribute to stress resistance, the underlying mechanisms for salt avoidance are unclear (Chen et al., 2021). Root halotropism is defined as a negative tropic response to high salinity (Galvan-Ampudia et al., 2013). Existing mutants defective in salt sensing or signaling did not show defective halotropic root bending, but instead showed enhanced bending (Figure 1), possibly due to higher Na^+^ accumulation in root tips, which might suggest that the ionic stress of Na^+^ triggers root halotropism. Besides ionic stress, high salinity also induces hyperosmotic stress. Central regulators of root hydrotropism, MIZ1 and cytokinin, did not contribute to root halotropism (Figure 2). However, ABA is shared by root halotropism and hydrotropism, downstream of salt and osmotic stress sensing (Dietrich et al., 2017; Miao et al., 2021) (Figure 3).

Based on our findings, we proposed a model in which the stress hormone ABA mediates root halotropism through phosphorylation regulation of the MT-associated protein SP2L (Figure 7). The gradually increasing salinity may induce ABA accumulation in root tips, which activates SnRK2 protein kinases. The activated SnRK2.6 interacts with and phosphorylates the MT-associated protein SP2L at Ser 406, which is essential for SP2L-mediated root halotropism (Figures 4 and 5). During halotropism, the initial transversely oriented cortical MTs reoriented to the longitudinal direction in approximately 2 hours in the elongating epidermal cells at the root transition zone, which requires SP2L function (Figures 6C and 6D). As the closest homolog of SPR2, SP2L has a similar role as SPR2 in mediating cortical MT array reorientation, together with the MT-severing enzyme katanin (Figures 6H-J). The cortical MTs serve as the trajectories for the CesA complexes and may mediate root halotropism by guiding cellulose microfibril patterns (Figures 6K-6Q and S4). Together with previous findings of the regulation of CesA complexes by cortical MTs (Endler et al., 2015; Gu et al., 2010), we outline a molecular mechanism for how the stress hormone ABA regulates the right-handed change in anisotropic cell expansion in elongating root cells during halotropism. However, it is unclear whether ABA-mediated SP2L phosphorylation is enough for anisotropic cell expansion and root halotropism. Recent studies reported that ABA promotes AHA2 phosphorylation and mediates an asymmetric H^+^ efflux required for differential growth in the elongation zone during hydrotropic root response (Dietrich et al., 2017; Miao et al., 2021). Whether this ABA-mediated asymmetric H^+^ efflux contributes to anisotropic cell expansion during root halotropism requires further studies.

**Figure 7.**
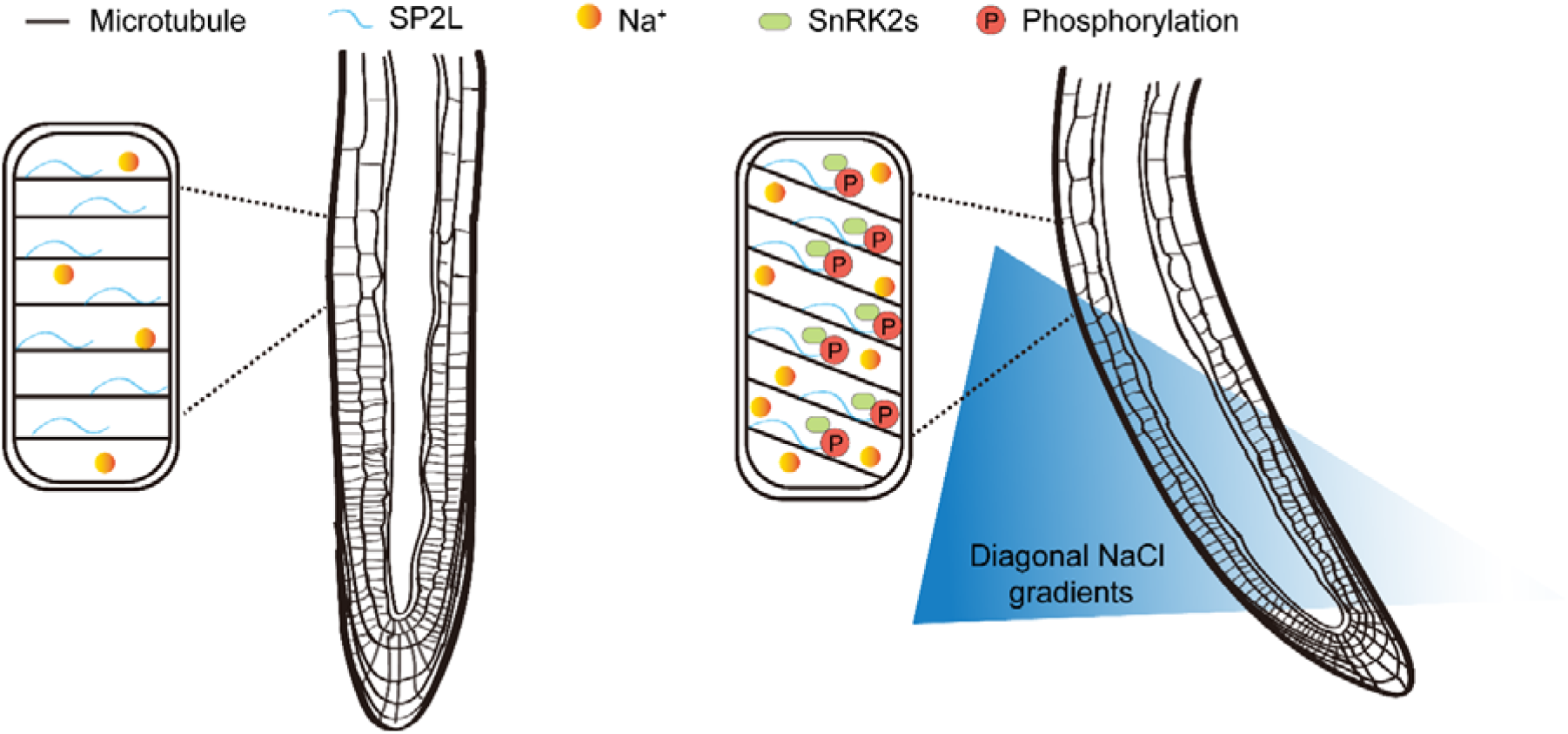
Schematic model for ABA- and SP2L-mediated root halotropism.

The halotropic root bending has to challenge or alter the root gravitropism, which would require auxin biosynthesis and redistribution. Based on our findings, mutants defected in gravitropism, including *aux1* and *pin2*, exhibited enhanced halotropic root bending (Figures 2A and 2B), indicated that gravitropism antagonizes halotropic root bending. The improved halotropic root bending in *aux1* and *pin2* may be caused by the damaged gravitropism but not enhanced halotropism. While PIN2 is a primary contributor to gravitropism, multiple PINs regulate auxin transport streams in the roots. We could not rule out the functional redundancy of these PIN proteins during halotropism. This also raises the question of how gravitropism is regulated during halotropic root bending. The auxin efflux carrier PIN2 undergoes quick phosphorylation and dephosphorylation regulation in response to environmental stimuli (Otvos et al., 2021; Wang et al., 2019; Yuan et al., 2020), but it remains unknown if SnRK2s phosphorylate PIN proteins to dampen gravitropism before root bending. Increased auxin levels were observed 2-6 hours at the side of the root facing the lower salt concentration, which is opposite to the auxin distribution for gravitropism (Galvan-Ampudia et al., 2013). Since auxin redistribution seems not to be required for halotropism based on mutant analysis in *Arabidopsis* (Figures 2A and 2B), it remains unclear if the aforementioned asymmetric auxin distribution challenges gravitropism to facilitate halotropic root bending. The right-handed cell elongation seems to be limited in the bending region, raising another question of how the change in anisotropic cell expansion is being recovered. This may be related to the dephosphorylation of SP2L or the re-establishment of root gravitropism. Indeed, ABA or salt stress-induced SnRK2 activation peaks at 0.5-1 hour and diminishes around 1.5-2 hours (Komatsu et al., 2013; Lin et al., 2020). The endocytosis of PIN2 occurs six hours after halo-stimulation (Galvan-Ampudia et al., 2013), which may somewhat dampen asymmetric auxin distribution (Retzer et al., 2019). Hence, it will be interesting to study the antagonistic regulations of tropisms during halotropic root bending.

Although auxin redistribution controls most tropic responses, it is not required for root hydrotropism and negative root phototropism in *Arabidopsis* (Shkolnik et al., 2016). Besides the widely accepted Cholodny-Went theory for tropic growth regulation, the asymmetric distribution of cytokinin confers an asymmetric cell division during hydrotropism. In addition to asymmetric cell elongation and division, we have revealed that a change in anisotropic cell expansion controlled by MT reorientation regulates a tropic response. This is consistent with earlier findings that phototropism requires blue light-induced MT array reorientation in elongating hypocotyl cells, which is controlled by the MT minus-end binding protein SPR2, the closest homolog of SP2L, together with the MT-severing enzyme katanin (Fan et al., 2018; Lindeboom et al., 2013; Nakamura et al., 2018). However, whether blue light-induced MT array reorientation induces a change in anisotropic cell expansion during phototropism requires further study. We found that SnRK2.6 also phosphorylates SPR2 at Ser414 (Figure S6), the conserved serine corresponding to Ser406 in SP2L. This phosphorylation could be detected in *Arabidopsis* using state-of-the-art mass spectrometry (Mergner et al., 2020). Whether this phosphorylation affects SPR2 function requires further investigation, especially in terms of phototropism. Further study is also needed to determine if any change in anisotropic cell expansion contributes to other tropisms such as phototropism and hydrotropism.

In summary, we have outlined several molecular mechanisms underlying root halotropism, in which the SnRK2-SP2L module controls a right-handed root twisting by mediating cortical MT reorientation and the microfibril deposition coordinate in root elongating cells. This indicates that anisotropic cell expansion mediates tropic responses where ABA, MT-reorientation and microfibril deposition have central roles.

## EXPERIMENTAL PROCEDURES

Further details and an outline of resources used in this work are provided in the Supplemental Experimental Procedures, including plant materials and plasmid constructs, vertical split-agar assay, visualization of Na^+^ using CoroNa Green, yeast two-hybrid assay, pull-down assay, bimolecular fluorescence complementation assay, IP-MS assay, *in vitro* phosphorylation assay, generation of anti-phosphorylation antibodies and immunoblotting, and microscopy.

## QUANTIFICATION AND STATISTICAL ANALYSIS

Student’s *t*-test and post-hoc Tukey test after ANOVA were used to determine the statistical significance between wild type and mutants in assays related to root curvature.

## SUPPLEMENTAL INFORMATION

Supplemental Information includes Supplemental Experimental Procedures and six figures.

## ACKNOWLEDGMENTS

This work was supported by the National Natural Science Foundation China (NSFC grant 31970293), the Strategic Priority Research Program of the Chinese Academy of Sciences (Grant No. XDB27040107), and the Shanghai Center for Plant Stress Biology, Chinese Academy of Sciences. S.P. acknowledges the financial aid of an ARC Discovery grant (DP19001941), Villum Investigator (Project ID: 25915), DNRF Chair (DNRF155) and Novo Nordisk Laureate (NNF19OC0056076), Novo Nordisk Emerging Investigator (NNF20OC0060564) and Lundbeck foundation (Experiment grant, R346-2020-1546) grants.

## AUTHOR CONTRIBUTIONS

Y.Z. conceived and designed the research. B.Y., W.Z., and L.X. performed the experiments. B.Y., W.Z., L.X., J.K.Z., S.P., and Y.Z. analyzed the results. Y.Z., B.Y. and S.P. wrote the manuscript.

