## Supplemental Data for "Root twisting drives halotropism via stress-induced microtubule reorientation"

**Supplemental Experimental Procedures**

**Plant materials**

All plants are in the Col-0 background unless indicated. Plants were grown at 23 °C under a 16/8 hr light/dark photoperiod with 60%–70% relative humidity.

The T-DNA insertion knockout mutant *sp2l-4* (SAIL_117_F12) was obtained from the Nottingham Arabidopsis Stock Center (NASC). The *SP2L* transcript in *sp2l-4* was analyzed using primers 5´-ATGCGATCACAAACCGCTTC and 5´-GAGATCCAACATTTTGACACGTAG. Professor Takashi Hashimoto provided the *sp2l-2* mutant (Yao et al., 2008).

Some plant materials used in this study include: *spr1-2* (N6547) (Nakajima et al., 2004), *spr2-2* (N6549) (Shoji et al., 2004), *mor1-1* (N67061) (Whittington et al., 2001), *mdp25* (SALK_022955) (Nagata et al., 2016), *vln4* (SALK_049058) (Zhang et al., 2011), *map65-1* (N67830) (Lucas et al., 2011), *afh14-1* (SALK_058886) (Li et al., 2010), and *ton1b* (SALK_087082).

The hydrotropism-related mutants *miz1-2* (SALK_076560) and *miz1-3* (SALK_126928) were obtained from the Nottingham Arabidopsis Stock Center (NASC) and have been reported (Kobayashi et al., 2007). The salt-sensing mutant *moca1* was provided by professor Zhenming Pei (Duke University) (Jiang et al., 2019). The salt stress-signaling fast neutron-mutagenized mutants *sos1-1* (N3862), *sos2-1* (N3863), *sos3-1* (N3864) were in Col *gl1* background(Liu et al., 2000; Liu and Zhu, 1998; Shi et al., 2000). Professor Jia Li provided the cytokinin-related mutants *ahk2-5/cre1-2*, *ahp1/2/3*, and *arr16/arr17* (Lanzhou University, China) (Chang et al., 2019). Professor Lin Xu provided the auxin-related mutants *pin2* (CS8058) (Roman et al., 1995), *aux1* (GABI_459H07)(Xu et al., 2007), *yuc9* (SAIL_871_G01), *pin1-11* (GABI_051A10), *pin7-2* (SALK_044687), *yuc1* (SALK_106293), *yuc2* (SALK_030199), *yuc4* (SM_3_16128), *yuc6* (SALK_093708), and *yuc1-D* (Cheng et al., 2006; Zhao et al., 2001) (Center of Excellence in Molecular Plant Sciences, Chinese Academy of Sciences). The ABA-related *snrk2.2/3/6* and *pyl* duodecuple mutants have been described previously (Fujii and Zhu, 2009; Zhao et al., 2018). The double mutant *nced3/5* was derived from a cross between *nced5‐2* (GK_328D05) and *nced3‐2* (GK_129B08) (Frey et al., 2012) and was obtained from professor Jing Zhang (China Agricultural University). The microfilament-related mutants *vln2* (SAIL_613_C03), *vln3*(SALK_078340), and *vln2/3* were obtained from professor Shanjin Huang (Tsinghua University) (Bao et al., 2012). The microtubule- and cellulose synthesis-related mutants, including *katanin1* (CS9531) (Wang et al., 2017), *cesa1* (*rsw1-1*, CS6554) (Arioli et al., 1998), *cesa3* (*ixr1-2*, CS6202) (Scheible et al., 2001), and *cesa6* (*prc1-1*, CS297) (Fagard et al., 2000), were obtained from the Nottingham Arabidopsis Stock Center (NASC) and have been reported. The ABA FRET sensors ABACUS1-2μ (CS68846) and ABACUS1-80μ (CS68847) have been reported (Jones et al., 2014). *csi1-4* (SALK_047252) and *csi1-5* (SALK_051146) was obtained from Alberto Macho (Shanghai Center for Plant Stress Biology).

**Growth Conditions**

The seeds were sterilized with 5% sodium hypochlorite for 10 minutes and washed with sterile-deionized water for 5 times. Seeds were grown vertically on 1.2% agar containing 1/2 MS nutrients (PhytoTech, M542), 1% sucrose, pH5.7. Plates were wrapped with tin foil and stratified at 4 ℃ for 3 days. For split-agar assay, 4-5 days vertical grown seedlings were transferred from 1/2 MS medium to split-agar medium with or without NaCl or mannitol. The seedlings grown on the medium were cultured in the Percival CU36L5 incubator at 23 ℃ under long-day conditions (16 h light, 22 °C; 8 h dark, 20 °C). Seedlings were grown in soil in a growth room at 22_C with a 65%–80% relative humidity under long-day conditions.

**METHOD DETAILS**

**Plasmid Construction and Plant Transformation**

To generate *SP2Lpro:SP2L* construct, the *SP2L* genomic fragment containing the 1617-bp *SP2L* promoter and the 537-bp 3'-UTR were amplified and cloned into the Gateway donor vector pENTR (Invitrogen), and then recombined into the Gateway destination vector pGWB513 using Gateway LR Clonase (Invitrogen). The non-phosphorylatable point mutation of *SP2L* (*SP2L^406A^*) was obtained by site-directed mutagenesis. The *SP2Lpro:SP2L* and *SP2Lpro:SP2L^406A^* constructs were confirmed by sequencing and transformed into the *sp2l-4* mutant using the *Agrobacterium*-mediated floral dip method (Zhang et al., 2006).

The *SP2L* and *SP2L^S406A^* CDS fragments were cloned into pCAMBIA1300 to generated *35S:SP2L-YFP* and *35S:SP2L^S406A^-YFP* constructs. The resultant plasmids were confirmed by sequencing and transformed into Col-0 background.

The *SP2L* CDS fragment was fused with *mCherry* and cloned into pCAMBIA1300 to generated the *35S:SP2L-mCherry* construct. The resultant plasmid was confirmed by sequencing and introduced into transgenic lines harboring *35S:GFP-TUB6*.

To generate the *pGBKT7-SP2L* and *pGADT7-SnRK2.6* constructs, the CDS fragments of *SP2L* and *SnRK2.6* were amplified and cloned into the *pGBKT7* and *pGADT7* vector.

To generate the *pGEX4T-1-GFP* constructs, the fragments of *GFP* was amplified and cloned into the *pGEX4T-1* vector.

To generate the *pGEX4T-1-SP2L-MC-GFP* and *pGEX4T-1-SP2L^S406A^-MC-GFP* constructs, the fragments of *SP2L-MC* (truncated CDS fragments of *SP2L, SP2LCDS^970-2463^*) and *SP2L^S406A^-MC* were amplified and cloned into the *pGEX4T-1-GFP* vector.

To generate the *pET-32a-SnRK2.6 and pMAL-c2X- SnRK2.6* constructs, the CDS fragments of *SnRK2.6* was amplified and cloned into the *pET-32a* and *pMAL-c2X* vector.

To generate the *pGEX6P-1-SPR2* constructs, the CDS fragments of *SPR2* was amplified and cloned into the *pGEX6P-1* vector.

**Vertical** **split-agar assay**

The vertical split-agar assay used in this study was modified from the reported method (Galvan-Ampudia et al., 2013). Briefly, the ½ MS medium containing indicated concentrations of NaCl or mannitol was poured into the 100 mm × 15 mm square disposable Petri dish with a handmade plastic separator (truncated ruler). After solidifying the medium, we removed the plastic divider and then poured the fresh ½ MS medium (or ½ MS supplemented with 1 μM oryzalin or 1 nM isoxaben) into the other side. Fresh split-agar medium was used immediately, and the seedling transferring would be completed within 2 hours. Usually, 10-12 seedlings were placed on a plate. We transferred the seedlings grown vertically on ½ MS medium for 4-5 days to the split-agar medium. The distance between the root tip and the salt/MS medium boundary was 0.3 mm. Subsequently, seedlings were grown vertically on the split-agar plate under same growth conditions. The photo was taken after halotropic treatment for 18-20h. ImageJ was used to analyze the root halotropic curvature (Schneider et al., 2012).

In the previous method (Galvan-Ampudia et al., 2013), seedlings were grown on 1/2 MS medium, and diagonal NaCl gradients were generated after 5 days post-germination without transferring seedlings. Thus, seed germination inconsistency leads to different root lengths, and the distance between root tips and the NaCl/MS interface varies greatly (Galvan-Ampudia et al., 2013). What we modified is that we selected and transferred the 4-5 day-old seedlings from 1/2 MS medium to the fresh diagonal NaCl gradients plate. Through selection, we ensured the consistency of seedling growth and accurately maintained the distance consistency between root tips and NaCl/MS interface. Our modification makes the halotropic treatment more stable and increases the number of effective samples per plate. Moreover, some mutants with germination defects and development defects can be selected to be comparable with wild type by transferring seedlings.

**Visualization of Na^+^ using CoroNa Green**

We transferred 4- to 5-day-old seedlings to a split-agar medium with 100 mM NaCl halostimulation for 3 h. After being rinsed three times in sterilized water, the seedlings were stained in 20 μM CoroNa Green (CoroNa™ Green Sodium Indicator, Thermo Fisher, C36676) for 1 hour, and then washed three times in sterilized water. The stained root tips were analyzed and imaged using a confocal microscope. The absorption and emission maxima of the CoroNa Green indicator are at approximately 492 and 516 nm, respectively. The fluorescence intensity was analyzed by ImageJ. The corona fluorescence intensity of Col-0 was normalized.

**Yeast two-hybrid assay**

The full-length CDS of *SnRK2.6* and *SP2L* were cloned into pGADT7 and pGBKT7, respectively. The resulting constructs were transformed into Y2H gold yeast strain (Takara Bio). Combinations of pGADT7-SnRK2.6 with pGBKT7 and pGADT7 with pGBKT7-SP2L were used as negative controls. Yeast transformants were selected on SD medium lacking Leu and Trp media and 5 μl of serial decimal dilutions transferred to SD medium lacking Leu, His, and Trp with 20 mg/ml X-α-Gal. Plates were kept at 28 ℃ for 3 to 4 days.

**Pull-down assay**

The pull-down assay was performed as described previously (Li et al., 2017). Purified GST-SP2L-MC-GFP or GST-GFP recombinant protein (5 μg) were conjugated on Glutathione-Sepharose and incubated with 5 μg ST-His-SnRK2.6 protein for 1 hour in 1 mL of incubation buffer (20 mM Tris-HCl, pH 7.4, 1 mM EDTA, and 100 mM NaCl). Then 50 μl of this 1-ml mixture was used as input. The bead-protein complex was rinsed three times with 1 mL of washing buffer (0.5% NP40, 1 mM EDTA, 20 mM Tris, pH 7.4, 300 mM NaCl), and then the proteins were eluted for immunoblot detection. The GST-GFP protein was used as a negative control.

**Bimolecular fluorescence complementation (BIFC) assay**

BiFC assays were performed using tobacco leaves (*Nicotiana benthamiana*) as described (Fang and Spector, 2010). The full-length CDS of *SnRK2.6* was fused with the N-terminal of YFP (amino acids 1-154) while *SP2L* was fused with YFPC (amino acids 155-238), respectively, and cloned into pCAMBIA1300. The resulting constructs were transformed into *Agrobacterium* strain GV3101. A single fresh colony was cultured and diluted to the absorbance at 600 nm (OD600) of ~1.0 and then injected into tobacco leaves. After 48 hours, the YFP signal was analyzed and imaged using a confocal microscope. The empty YFPN and YFPC plasmids were used as negative controls.

**IP-MS Assay**

Five-day-old transgenic seedlings were ground in extraction buffer [50 mM Tris-HCl, pH 7.5, 150 mM NaCl, 5 mM MgCl_2,_ 0.5 mM DTT, 10% glycerol, 0.1% NP-40, 1×Protease and Phosphatase Inhibitor Cocktail (Thermo Fisher, Cat#: 78442), 1 mM PMSF]. After being centrifuged at 4 ℃ for 30 min at 17115 × g (13,500 rpm), the supernatants were incubated with anti-GFP magnetic agarose beads at 4 ℃ for 3 hours. After incubation, the anti-GFP magnetic agarose beads were washed two times with protein extraction buffer and one time with 1×PBS. The magnetic beads were retained in 100 μl 1×PBS for mass spectrum identification (Plant Proteomics and Metabolomics Core Facility, Shanghai Center for Plant Stress Biology, CAS).

***In vitro* phosphorylation assay**

The *in vitro* phosphorylation assay was performed as previously described (Zhao et al., 2018). Briefly, 0.1 μg purified ST-His-SnRK2.6 recombinant protein was incubated with 1 μg purified GST-SP2L-MC-GFP or GST-GFP recombinant protein for one hour at 30 ℃ in reaction buffer (25 mM Tris-HCl, pH 7.5, 10 mM MgCl_2,_ 0.25 mM DTT, 1 mM ATP, 2 μl Ci [γ-^32^P] ATP). After incubation, proteins were separated by 12.5% SDS-PAGE gel and stained with Coomassie blue, then dried under vacuum at 80 °C for 1.5 h on filter paper. Phosphorylation was visualized by autoradiography.

**Generation of anti-phosphorylation antibodies and immunoblotting**

Anti-phosphorylation antibodies were generated in rabbit by ABclonal Technology (China). The HPLC high purity modified peptide C-KLEKRG-pS-GD was used as an antigen to generate polyclonal anti-phosphorylation antibodies for Ser406 in SP2L. The phosphorylation site-specific antibodies were purified using the non-modified peptide C-KLEKRGSGD, which absorbed the non-specific antibodies.

Total proteins were extracted in 5-day-old *35S*:*SP2L-YFP* and *35S*:*SP2L^S406A^-YFP* transgenic plants treated with or without 200 mM NaCl or 100 μM ABA. Total proteins were extracted in extraction buffer as stated above, separated by 12.5% SDS-PAGE, and then transferred to PVDF membrane for immunoblot detection. Western blot analyses were performed as previously described (Zhao et al., 2018). Anti-GFP antibody was used as a control.

**Microscopy**

Four-day-old seedlings expressing YFP-NPSN12 (Geldner et al., 2009) were transferred to a split-agar medium, then observed by a 20× objective of a spinning-disk confocal microscope equipped with a vertical stage for 2 h. The spinning-disk confocal microscopy is an inverted Nikon Ti-E microscope with a CSU-W1 spinning disk head (Yokogawa, Japan) with a deep-cooled evolve charge-coupled iXon Ultra 888 EM-CCD camera (Photometrics Technology, USA), and acquired by MetaMorph software (Molecular Devices). YFP signal was imaged using a 514-nm laser and a 540/50-nm emission filter.

To observe the reorientation of microtubules, we put seeds on the right side of the 1/2 MS medium in chambered cover slides (ibidi, 80287) and pushed the seeds close to the cover slide. After growing vertically for 4 days at 23°C, the chambered cover slides were used for imaging directly. With halo-stimulation, we replaced the left corner of the 1/2 MS medium with 1/2 MS medium with 200 mM NaCl. Seedlings stayed upright during this process.

Cortical microtubules and CesA3 were observed by a 100× objective (Apo TIRF, NA 1.49) on a vertical converter of an inverted Nikon Ti2-E spinning disk confocal microscope equipped with CSU-W1 spinning disk head (Yokogawa, Japan), iXon Life 888 EM-CCD and acquired by 3i Slidebook software (DSS Imagetech). GFP and mCherry signals were detected with a 488-nm laser and a 482/35-nm emission filter and a 561-nm laser and a 542/27-nm emission filter, respectively.

**ABA dynamics detection**

Details of the method for detecting abscisic acid dynamics in roots by FRET sensor have been described previously (Jones et al., 2014; Xie et al., 2020). Root ABA dynamic detection was performed after 0 and 2 hours of halostimulation. Confocal photos were taken by Olympus FV3000 with a 20× objective. The excitation of donor (CFP) and receptor (YFP) were 442 and 514 nm, respectively. The donor fluorescence emission is set at 458–482 nm, and the acceptor fluorescence emission is set at 520–550 nm. Fluorescence resonance energy transfer from donor to acceptor was measured under 442 nm excitation (CFP) and 520–550 nm emission (YFP). The ratio of FRET acceptor emission (Am) to FRET donor emission (Dm) after excitation of the FRET donor (Dx) was calculated as an approximation of FRET ratio. DxAm/DxDm data were normalized to control. DxAm and DxDm intensity values were measured for a defined ROI and the background fluorescence in a region outside of the root elongation zoon was subtracted. Image processing and analysis were performed using ImageJ (Schneider et al., 2012). YFP is divided by CFP background to obtain FRET images. Images were displayed in 16 colors.

**Root gravitropic bending assay**

The seedlings grown vertically for 5 days were transferred to 1 / 2 MS medium, and the plates were immediately turned 90 degrees compared with the original vertical position. After 24 h of the gravistimulation, the root bending angle was measured by Image J.

**Supplemental Figures**

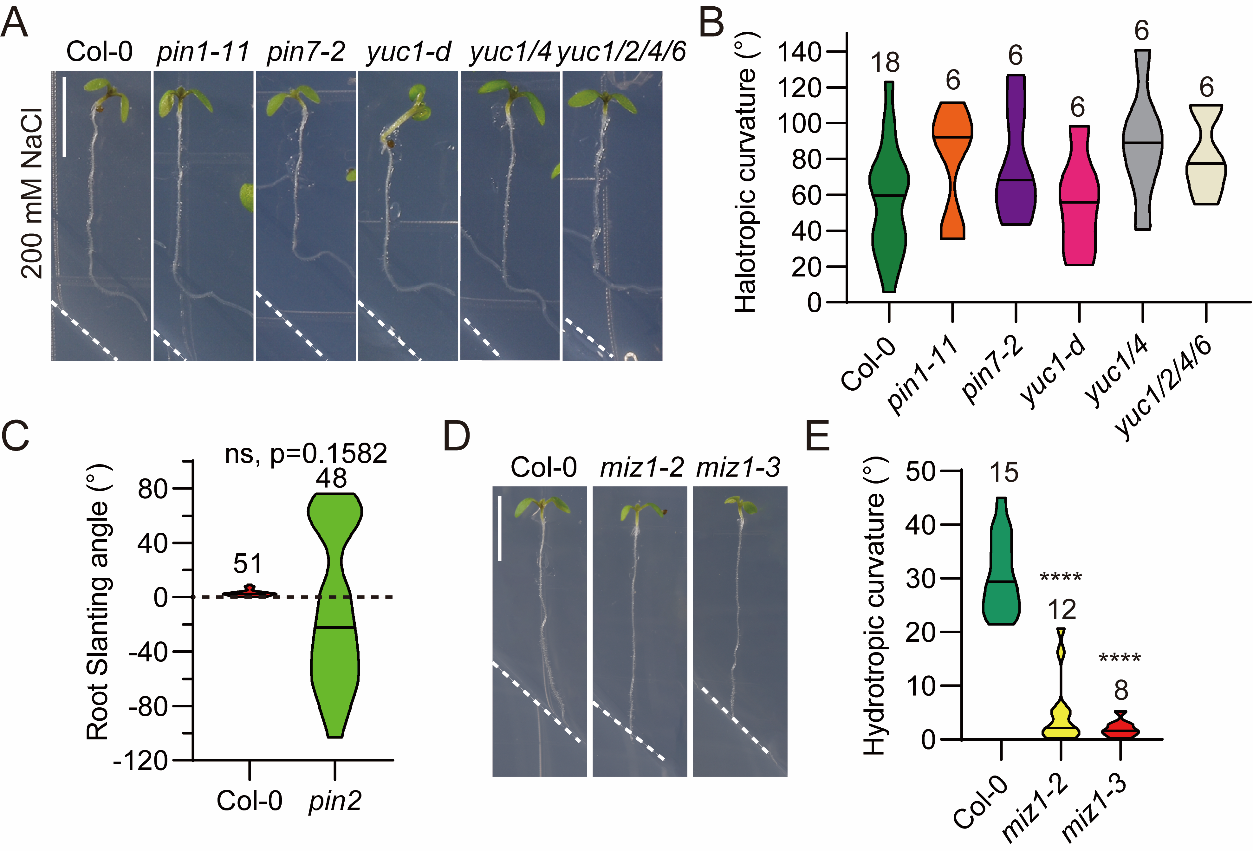

**Figure S1. Halotropic and hydrotropic root bending in auxin-related mutants and ER Ca^2+^-ATPase inhibitor** ***miz1* mutants. Related to Figure 2.**

(A-B) Root halotropic growth of Col-0 wild-type and auxin biosynthesis and transport mutants (A). The halotropic root curvature was quantified (B). Scale bar, 0.5 cm.

(C) Wild-type (Col-0) and *pin2* mutant seedlings were transferred to the control split-agar medium without NaCl, and the halotropic root curvature was quantified.

(D-E) Root hydrotropic growth of Col-0 wild-type and *miz1* mutants (D). The hydrotropic root curvature was quantified (E). **** *P* <0.0001, Student’s *t*-test. Scale bar, 0.5 cm.

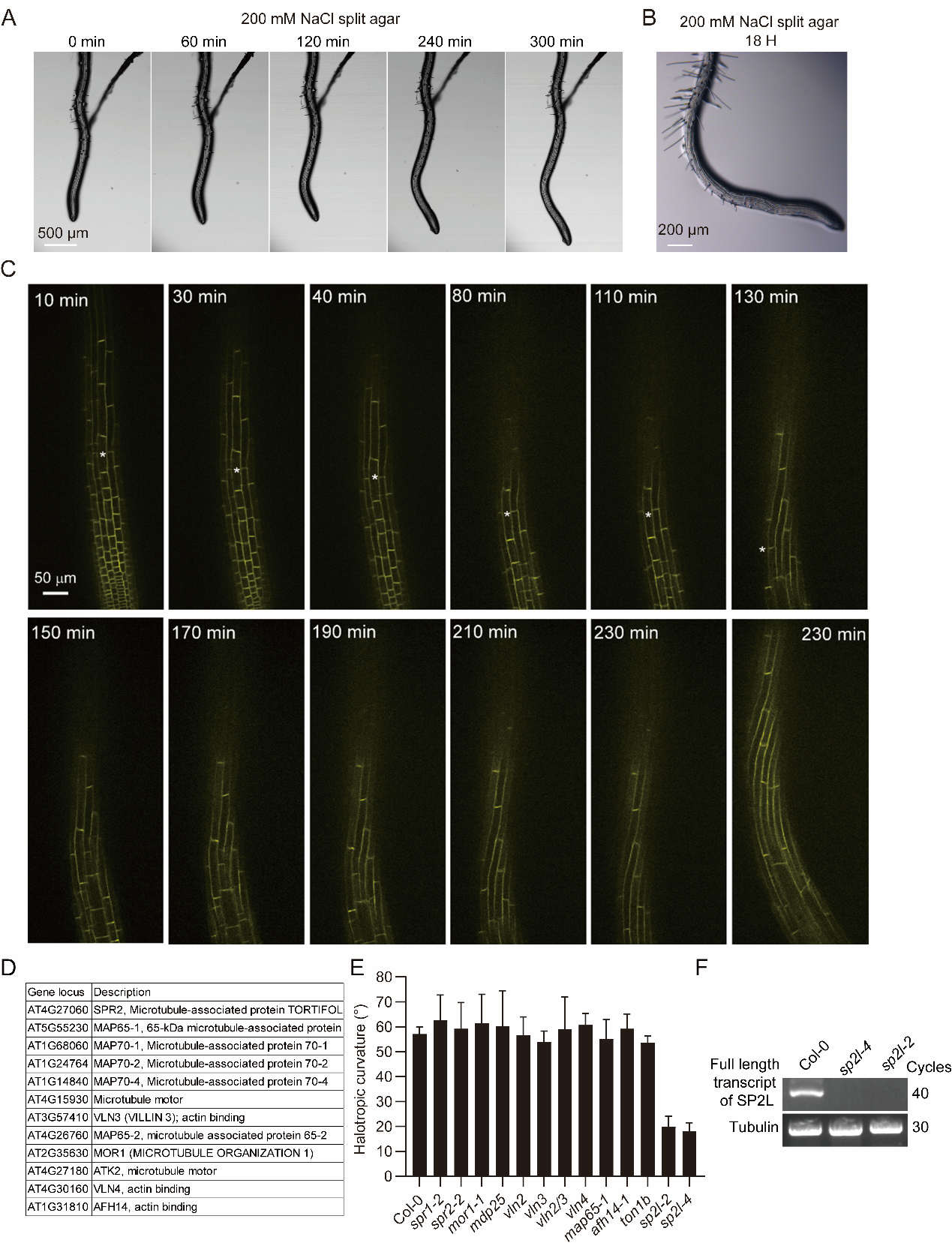

**Figure S2. Halotropic root growth in *Arabidopsis* Col-0 wild-type seedlings and mutants for putative SnRK2 cytoskeleton-related substrates*.* Related to Figure 3.**

(A-C) Four-day-old Col-0 wild-type seedlings with straight roots were transferred from ½ MS vertical plates to halostimulating ½ MS split-agar medium containing 200 mM NaCl at the bottom left side. Root growth was monitored using wild-type seedlings at the corresponding time points under a microscope (A), stereoscope (B), and a Nikon-Andor WD spinning disc confocal with a vertical stage (C).

(D) Cytoskeleton-related proteins that may be putative SnRK2 substrates.

(E) Halotropic root curvature was quantified in mutants of cytoskeleton-related genes. (F) RT-PCR analysis of *sp2l-4* and *sp2l-2* mutants. Tubulin was used as a loading control.

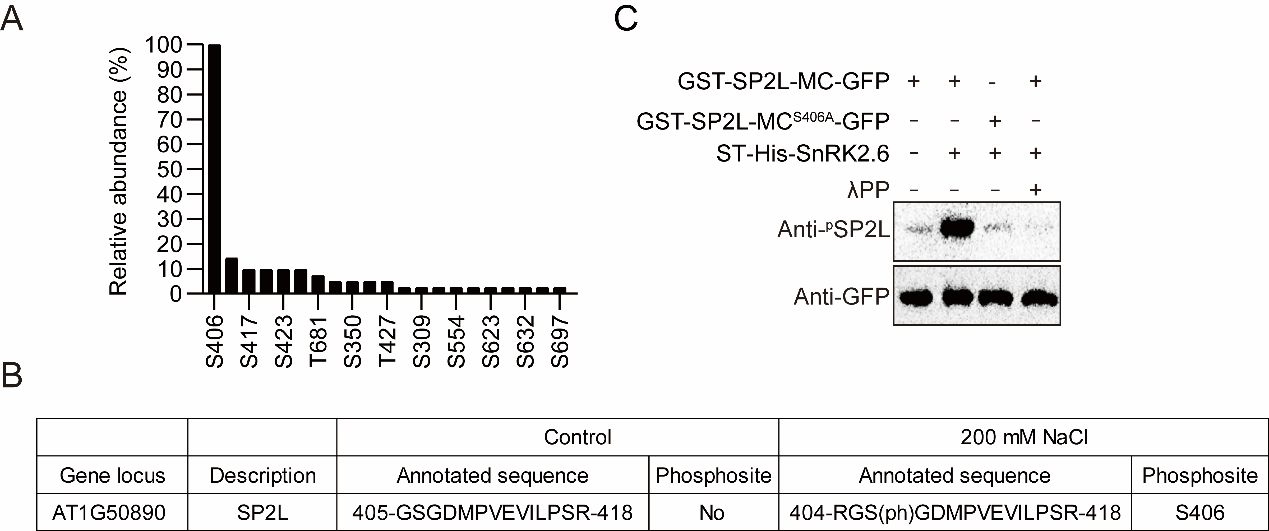

**Figure S3. SnRK2.6 phosphorylates SP2L at Ser406. Related to Figure 4.**

(A) The relative abundance of phosphosites detected by mass spectrometry after *in vitro* phosphorylation assay. The most abundant phosphosite (S406) was set to 100%. S: serine; T: threonine.

(B) Mass spectrometry identified a salt-induced SP2L phosphorylation site *in vivo.*

(C) Phosphorylation of wild-type and Ser-to-Ala mutated GST-SP2L-MC-GFP by recombinant ST-His-SnRK2.6. Anti-phospho-Ser406-SP2L antibodies were used to detect phosphorylation of SP2L. GFP was used as a loading control. λPP: Lambda protein phosphatase.

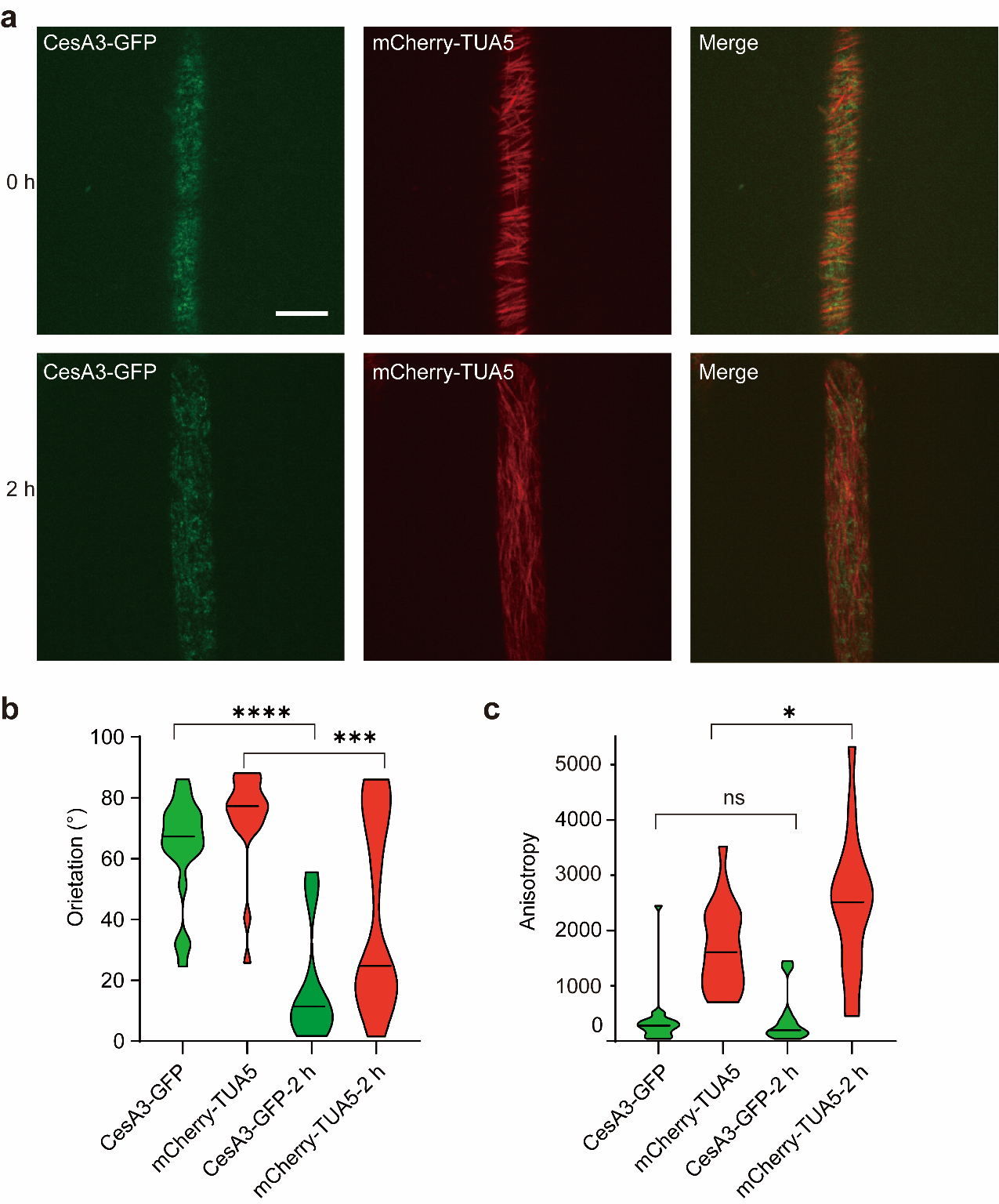

**Figure S4. Reorientation of cellulose microfibril patterns upon halo-stimulation. Related to Figure 6.**

(A) CesA3-GFP (left) and mCherry-TUA5 labeled MTs (middle) in root epidermal cells in the transition zone after 2 hours of halostimulation. The merged image is shown on the right. Scale bars, 10 μm.

(B-C) Quantification of CesA3-GFP and cortical MT orientation and anisotropy in (A). Data are mean values from 18-20 independent cells.

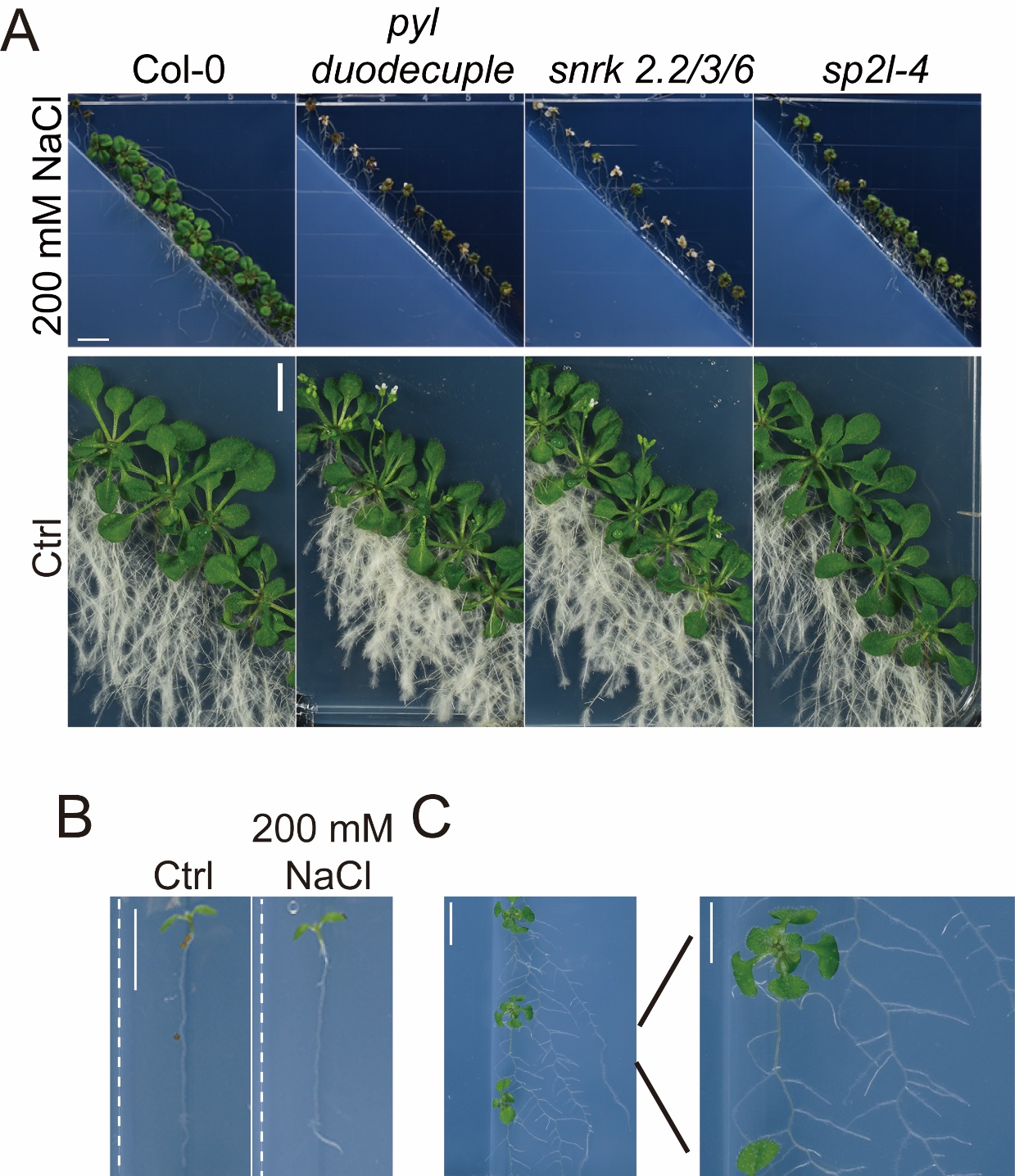

**Figure S5. Halotropism contributes to salt stress tolerance or avoidance. Related to Figure 7.**

(A) Seedling growth of Col-0 wild-type, ABA signaling mutants (*pyl* duodecuple, *snrk2.2/3/6)*, and *sp2l-4* mutant 21 days after transferring seedlings to a split-agar medium with 200 mM NaCl (up panel) and control split-agar medium (down panel). Scale bars, 1 cm.

(B) Wild-type (Col-0) seedlings were transferred to split-agar medium with 200 mM NaCl that generates vertical NaCl gradients. Scale bars, 0.25 cm.

(C) Scale bar, 0.5 cm.

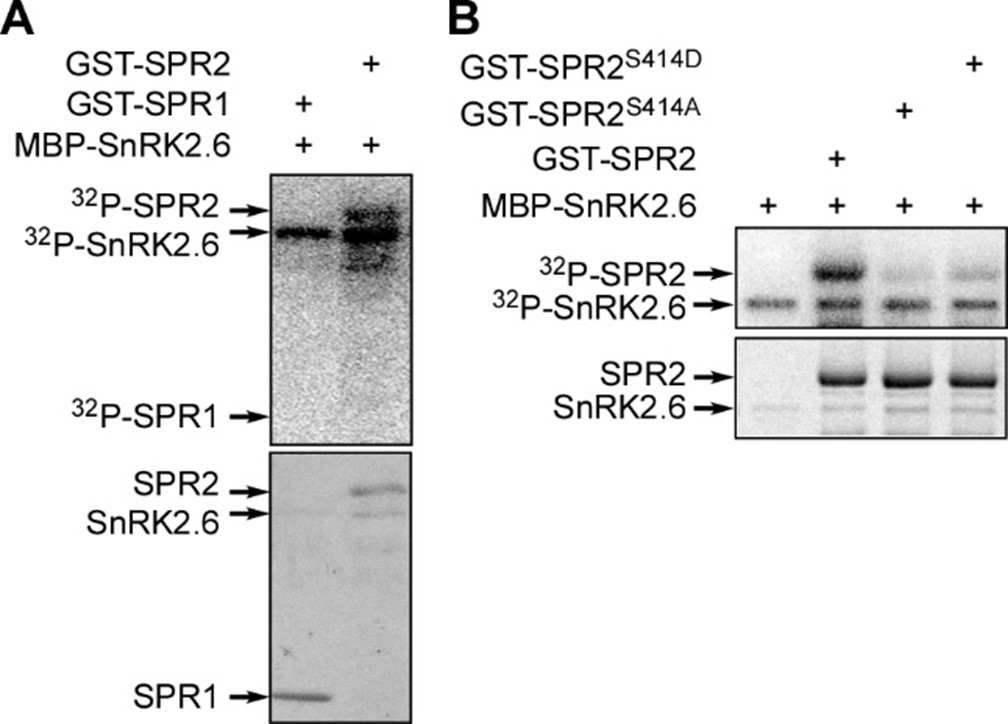

**Figure S6. SnRK2.6 phosphorylates SPR2 at Ser414. Related to Figure 7.**

Phosphorylation of wild-type (A) and Ser-to-Ala (B) mutated full-length GST-tagged SPR2 by recombinant MBP-SnRK2.6. GST-SPR1 was used as a negative control. Autoradiography (upper panel) and Coomassie staining (lower panel) exhibit phosphorylation and loading of proteins.
